# Mannose Supplementation Curbs Liver Steatosis and Fibrosis in Murine MASH by Inhibiting Fructose Metabolism

**DOI:** 10.1101/2024.01.17.576067

**Authors:** John G. Hong, Yvette Carbajal, Joshaya Trotman, Mariel Glass, Victoria Sclar, Isaac L. Alter, Peng Zhang, Liheng Wang, Li Chen, Matthieu Petitjean, Scott L. Friedman, Charles DeRossi, Jaime Chu

## Abstract

Metabolic dysfunction-associated steatohepatitis (MASH) can progress to cirrhosis and liver cancer. There are no approved medical therapies to prevent or reverse disease progression. Fructose and its metabolism in the liver play integral roles in MASH pathogenesis and progression. Here we focus on mannose, a simple sugar, which dampens hepatic stellate cell activation and mitigates alcoholic liver disease *in vitro* and *in vivo*. In the well-validated FAT-MASH murine model, oral mannose supplementation improved both liver steatosis and fibrosis at low and high doses, whether administered either at the onset of the model (“Prevention”) or at week 6 of the 12-week MASH regimen (“Reversal”). The*in vivo* anti-fibrotic effects of mannose supplementation were validated in a second model of carbon tetrachloride-induced liver fibrosis. *In vitro* human and mouse primary hepatocytes revealed that the anti-steatotic effects of mannose are dependent on the presence of fructose, which attenuates expression of ketohexokinase (KHK), the main enzyme in fructolysis. KHK is decreased with mannose supplementation *in vivo* and *in vitro,* and overexpression of KHK abrogated the anti-steatotic effects of mannose. Our study identifies mannose as a simple, novel therapeutic candidate for MASH that mitigates metabolic dysregulation and exerts anti-fibrotic effects.

## Introduction

Metabolic dysfunction-associated steatotic liver disease (MASLD)^1^ is the most common liver disease worldwide. MASLD progression from simple steatosis to steatohepatitis (metabolic dysfunction-associated steatohepatitis or MASH) can lead to cirrhosis and liver cancer, necessitating liver transplantation^2,3^. Despite its prevalence, there are currently no approved therapies to prevent or reverse the progression of MASH. Therapeutic approaches have largely been siloed to either target metabolic dysregulation and cell injury or focus on inflammation and fibrosis^4^.

Fructose plays a key pathogenic role in the progression of MASH^5–7^. Excess fructose intake induces *de novo* lipogenesis (DNL), decreases beta-oxidation, and induces oxidative stress in the endoplasmic reticulum (ER) and mitochondria in hepatocytes^5–7^. Limiting fructose intake has been effective in clinical trials. Limiting dietary sugars reduces DNL and hepatic steatosis in children, and there is a direct relationship between MASLD risk and fructose intake in adults^8–11^. Therefore, targeting intracellular fructose metabolism is of particular interest^12^.

Ketohexokinase (KHK) is a key target in fructose metabolism by catalyzing the first committed step in fructose metabolism: Fructose➔ Fructose 1-phosphate. KHK has been positively linked to MASLD progression; KHK is increased in mouse models of MASLD^13,14^, and patients^14,15^. Increased KHK provokes ER stress in hepatocytes^14,16^. In pre-clinical studies, inhibition of KHK was effective in mitigating MASH steatosis and fibrosis in mouse models^14,17–19^. Recently, phase 2 studies using a KHK inhibitor in adults with MASLD have demonstrated a reduction in hepatic steatosis with appropriate safety and tolerability (NCT03256526^17^, NCT03969719^18^); KHK is also a target for ongoing phase 2 trials in MASLD lipid accumulation and insulin sensitivity (NCT05463575). However, the effect of KHK on liver fibrosis has not been evaluated.

Mannose is a simple sugar and C _2_ epimer of glucose. Its therapeutic potential has been long overlooked. Oral mannose supplementation is well-tolerated, readily accessible, and has completed a phase 3 trial for urinary tract infections (NCT01808755)^20^. Recently, mannose supplementation has emerged to play disease-modifying roles in obesity^21^, diabetes^21,22^, cancer^11,23,24^ and osteoarthritis^25^. Our group has previously shown that mannose supplementation can dampen hepatic stellate cell (HSC) activation*in vitro* and decrease fibrogenesis *in vivo*^26^. Hu et al. demonstrated an anti-steatotic effect of mannose in a mouse model of alcoholic liver disease^27^. Given the individual patient and public health burdens of MASLD and the scarcity of therapeutic options, we sought to determine whether mannose would have therapeutic benefits for MASH.

To study the therapeutic role of mannose in MASH, we used the well-validated Fat-and-Tumor MASH (FAT-MASH) model^28–31^ which consists of a 12-week high-fat, high-fructose diet with once weekly, low dose carbon tetrachloride (CCl_4_) intraperitoneal injections; this regimen has been shown to recapitulate the steatosis and fibrosis seen with advanced MASH in humans with MASH-fibrosis by 6 weeks^32^. In this study, we tested whether mannose supplementation could: 1) prevent MASH development when given concurrently with the 12-week MASH regimen, and 2) reverse established steatosis and fibrosis when started at 6 weeks in the setting of already-established MASH. High resolution digital pathology and Artificial Intelligence (AI)^32,33^ were used to quantify the histological phenotypes of steatosis and fibrosis and the related intervention effects. Mannose, in both the prevention and reversal groups, displayed anti-steatotic and anti-fibrotic effects in the FAT-MASH model. Transcriptomic analysis of FAT-MASH mouse livers and *in vitro* hepatocyte lines treated with mannose revealed dampening of fructose uptake and metabolism pathways. Specifically, we found that mannose dampened KHK, which led to decreased steatosis. KHK overexpression abrogated the anti-steatotic effect, demonstrating that mannose acts via KHK regulation. Together, these data point to a novel therapeutic role for mannose in the treatment of MASH.

## Results

### Mannose attenuates and reverses steatosis in the FAT-MASH mouse model

Mannose has been reported to ameliorate hepatic steatosis in alcoholic liver disease^27^, but whether mannose is efficacious in reducing steatosis in MASH is unknown. To examine mannose as a potential therapy for MASH *in vivo*, we used the well-validated FAT-MASH mouse model^28–31^. Eight-week-old male C57BL/6J mice were given a high fat, high cholesterol, high sugar diet with low dose CCl_4_ for 12 weeks, and placed *ad libitum* on either a low (5% weight/volume) or high (20% weight/volume) dose of mannose in the drinking water (Figure 1A, Materials and Methods). Furthermore, mannose supplements were initiated at either the start of the protocol (“prevention” group) or at 6-weeks into the 12-week FAT-MASH regimen, when steatosis and fibrosis are already present (“reversal” group). All mice gained weight similarly for the duration of experiments, regardless of diet or mannose supplementation group (Figure S1A).

**Figure 1.**
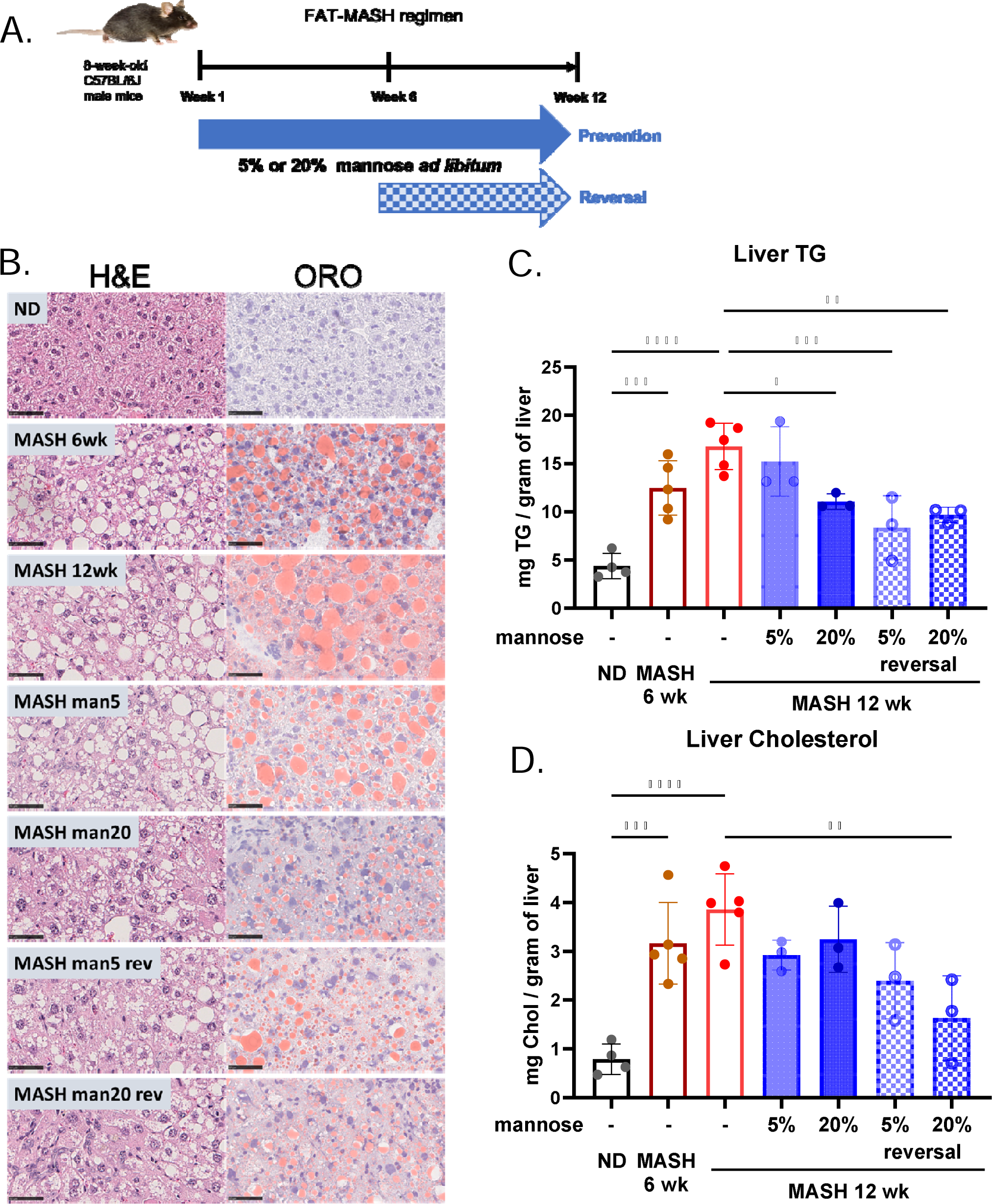
Mannose supplementation in FAT-MASH mice alleviates steatosis and lipid profile. (A) Graphical outline of the FAT-MASH regime. Mice were treated with ND or western diet with low dose CCl_4_ (MASH) for 12 weeks, without or with 5% or 20% mannose ad libitum. (B) Representative images of H&E and ORO staining from whole liver sections. Scale bar: 50µm. (C-D) Lipid profile of Hepatic TG (C) and cholesterol levels (D) in whole liver tissue (n=3-5). Results are expressed as mean ± SD with analysis by one-way ANOVA with Šidák post hoc test (*p<0.05, **p<0.01, ***p<0.001, and ****p<0.0001).

Consistent with previous reports^31,34^, mice that underwent FAT-MASH regimen for 12 weeks had histological evidence of MASH with extensive hepatic steatosis, along with elevated liver triglycerides (TG), liver cholesterol, liver weight, and liver/body weight ratio compared to normal diet (ND) (Figures 1B-D, S1A-C). Some features of MASH were already present by 6 weeks of the FAT-MASH regimen, including hepatic steatosis, elevated liver TG and cholesterol (Figures 1B-D). We measured serum alanine aminotransferase (ALT), aspartate aminotransferase (AST), and total and direct bilirubin in all groups. AST was significantly elevated and ALT trended higher in FAT-MASH mice after 12 weeks (Figures S1D-E). Mannose supplementation decreased mean AST and ALT values when compared to MASH, but only met statistical significance in AST with low dose mannose treatment in the reversal group. Serum total and direct bilirubin remained in the normal range across all groups at 12 weeks (Figures S1F-G).

Both low and high dose mannose supplements had a clear effect on alleviating steatosis, qualitatively observed by hematoxylin and eosin (H&E) and Oil Red O (ORO) staining with noticeably reduced size and number of lipid vesicles (Figure 1B). There was a dose-dependent effect of mannose treatment in both prevention and reversal groups (Figure 1B). Consistent with qualitative observations, mannose decreased levels of liver TG and cholesterol. Mannose reduced liver TG in all groups and met statistical significance except for low dose treatment in the prevention group (Figure 1C). Similarly, mannose decreased liver cholesterol in all groups with high dose treatment in the reversal group reaching statistical significance (Figure 1D). Liver weight and liver/body weight ratios were significantly reduced with high dose mannose treatment in both prevention and reversal groups (Figure S1B-C). Mean serum AST and ALT were decreased with mannose treatments, with low dose mannose treatment in the reversal group reaching statistical significance when compared to FAT-MASH mice without mannose treatment (Figure S1D-E). Mannose had no effect on serum AST and ALT with high dose mannose treatment in the reversal group. Total and direct serum bilirubin were reduced with high dose mannose treatment in the prevention group (Figures S1F-G).

To quantify the magnitude of steatosis, we applied a high resolution automated single fiber, single vacuole quantitative AI-based platform (FibroNest^TM^, Princeton, NJ, USA). This platform quantifies multiple phenotypic traits that account for fibrotic and steatotic severity and abundance^32,33^. Consistent with the qualitative histopathology observations (Figure 1B), FibroNest^TM^ quantitative measurements confirmed that the FAT-MASH regimen induced steatosis by 6 weeks and peaked at 12 weeks. There were significant increases in macro-steatosis area from ND at 0.3% to 12-week FAT-MASH regimen at 24.0 % (Figure 2B, Supplemental Table 1). A substantial increase in fat vacuole counts was also observed in 12-week FAT-MASH regimen when compared to ND (Figure 2C). Mannose alleviated the severity of steatosis, indicated by reduced size of steatosis area and abundance of steatotic vesicles using fat vacuole counts (Figures 2A-C). The anti-steatotic effect was apparent for both medium (6-18 um) and large (> 18 um) vacuoles, with a more robust effect on large vacuoles (Figure 2B). High dose mannose treatment was more effective than low dose mannose in both prevention and reversal groups (Figures 2A-C). Low and high dose mannose treatments in the prevention group reduced steatosis by 26% and 60% respectively, with high dose mannose reaching statistical significance. Similar reductions in steatosis were seen in the reversal group; low and high dose mannose treatments reduced steatosis by 43% and 70% respectively (Figure 2C, Supplemental Table 1).

**Figure 2.**
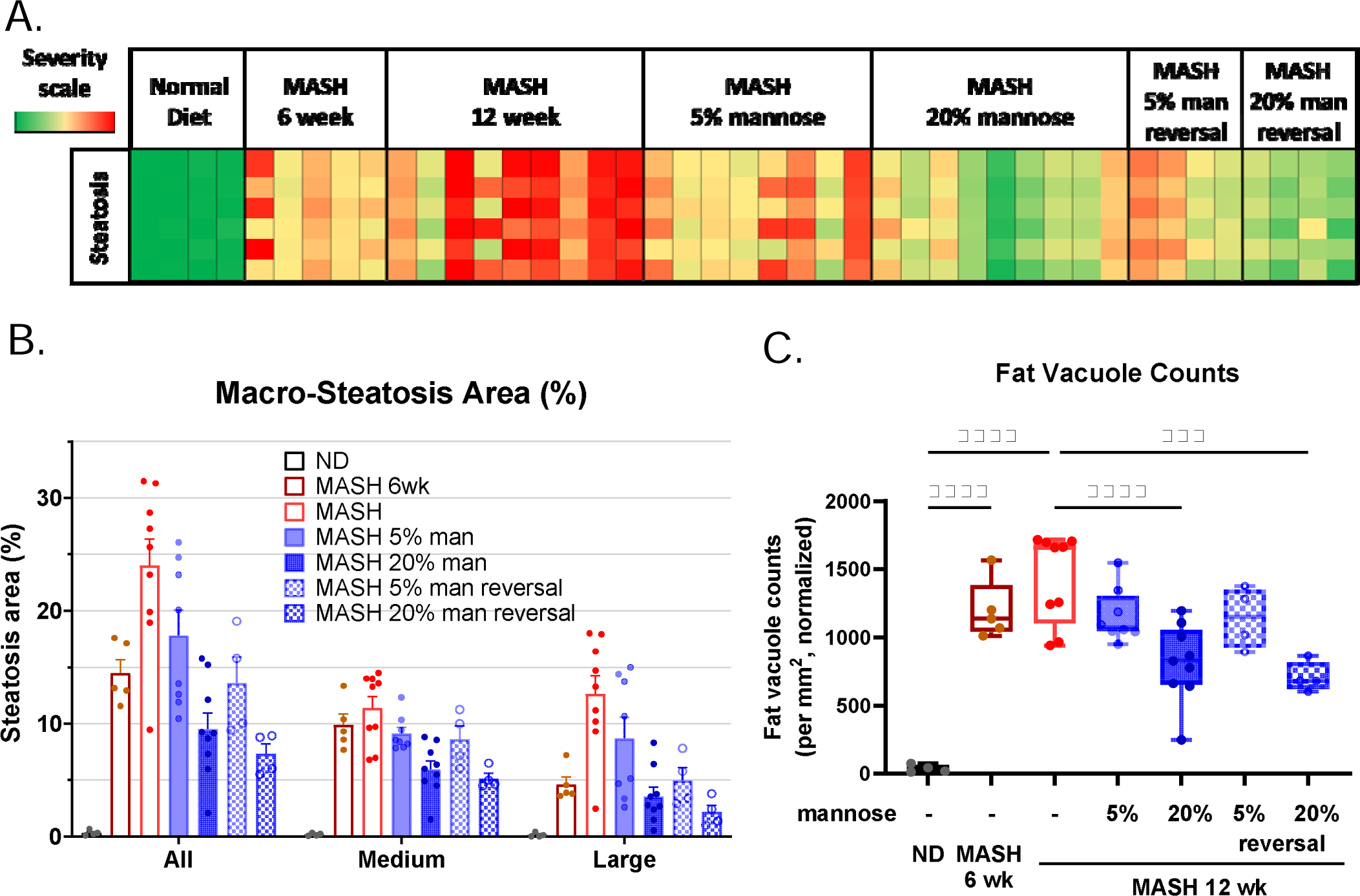
AI quantification of steatosis in FAT-MASH mice. (A) AI-based quantification of steatosis (Pharmanest) in mouse livers from treatment groups, with increasing severity indicated by heat chart color (green ➔ red). Columns are for individual mice, with n=4-9 for each treatment group. (B) Bar chart depicting quantification of steatosis area (%), for all, medium (6-18 um) or large (>18 um) vesicles. Individual mice are indicated by points, and statistical significance shown in Table S1. (C) Box plot showing total number of fat vacuoles for each treatment group (n=4-9) and compared by Student’s t-test (***p<0.001, and ****p<0.0001). ND, normal diet; MASH, FAT-MASH diet; man, mannose

Together, these data demonstrate that oral mannose supplementation in the FAT-MASH mouse model dampens liver steatosis, liver TG, and liver cholesterol, and points to mannose supplementation as a novel therapeutic for MASH-associated steatosis with the potential to both prevent and reverse steatosis.

### Mannose mitigates and reverses MASH fibrosis

The degree of fibrosis associated with MASH is an important driver of patient outcomes^35^. We have previously shown that mannose supplementation *in vitro* can mitigate the activation of HSCs, the major cell type responsible for liver fibrogenesis^26^, but the *in vivo* effects of mannose on fibrosis in MASH have not yet been studied. We therefore examined if mannose supplementation could reduce the severity of MASH fibrosis in the FAT-MASH mouse model. Consistent with previous reports^31,32,34^ mice fed the FAT-MASH diet developed substantial bridging fibrosis by 12-weeks, qualitatively shown with Sirius red staining for collagen content (Figure 3A). Importantly, both low and high dose mannose supplementation effectively mitigated liver fibrosis in both prevention and reversal treatment groups (Figure 3A). Western blots on whole liver lysates confirmed increased collagen (COL1A1) in FAT-MASH livers when compared to ND (Figure 3B). COL1A1 was decreased with low and high mannose treatments in the prevention group (Figure 3B).

**Figure 3.**
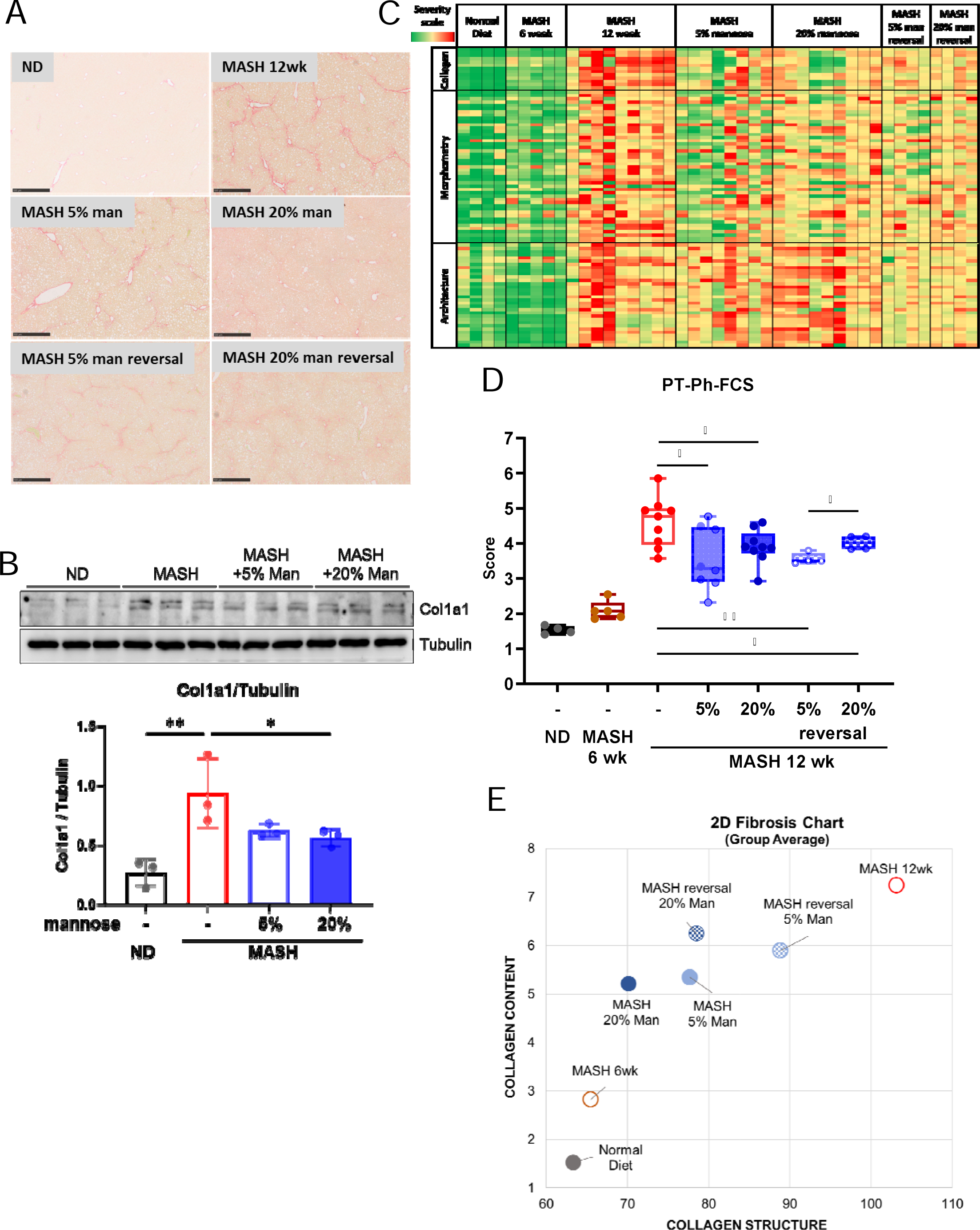
Mannose attenuates fibrosis in FAT-MASH mice. (A) Representative images of Sirius Red staining for collagen from whole liver sections. Scale bar: 500µm. (B) Western blot analysis of Col1a1 in whole liver lysates from ND or MASH mice without or with mannose supplements. Tubulin is used as a loading control, and Col1a1/Tubulin ratios are quantified with ImageJ, and compared by one-way ANOVA with Dunnett’s post hoc test for multiple comparisons (*p<0.05, **p<0.01). (C) AI-based quantification of Sirius red collagen staining. (D) Principal qFTs are aggregated into a Phenotypic Fibrosis Composite Score (PT-Ph-FCS). (E) 2D Fibrosis Chart shows the relationship between the collagen deposition and its structure. n=4-9 mice per group. Results expressed as mean ± SD and compared by Student’s t-test (*p<0.05, **p<0.01, ***p<0.001, and ****p<0.0001). ND, normal diet; MASH, FAT-MASH diet; man, mannose

To obtain an unbiased and comprehensive analysis of fibrosis in FAT-MASH mouse livers, we utilized the FibroNest^TM^ quantitative AI approach. FibroNest^TM^ measured three layers of fibrosis subphenotypes including: 1) collagen deposition (quantity), 2) morphometry (fiber shape and size), and 3) fibrosis architecture (fiber organization and complexity). The principal qFTs (quantitative Fibrosis Traits) for these 3 fibrosis subphenotypes are displayed in a phenotypic heat map from green to red, with red representing the most severe fibrosis (Figure 3C). The ND livers had the least fibrosis and the MASH livers at 12 weeks had the most severe fibrosis. Fibrosis severity was reduced in every mannose treatment group (Figure 3C). The principal qFTs were combined into a continuous Phenotypic Fibrosis Composite Score (PT-Ph-FCS), which normalizes to the parenchymal tissue present. Similarly, all groups receiving mannose treatment showed a significant reduction in the PT-Ph-FCS as compared to the MASH (12 weeks) (Figure 3D, Supplemental Table 2). In a 2D Fibrosis Chart of histological fibrosis traits, all mannose treatments in both prevention and reversal groups restored the collagen profile closer to the normal diet group (Figure 3E). Thus, mannose supplementation at low and high dose, and in prevention and reversal groups, all ameliorated fibrosis severity in MASH. Given that mannose supplementation directly mitigates HSC activation ^26^, we hypothesized that mannose has direct anti-fibrotic effects, independent of steatosis. To test this, we used the chemotoxic CCl_4_ mouse model^36,37^ to induce pure fibrosis with no steatosis. We used a similar treatment regimen (low and high dose mannose, under prevention and reversal timelines) to examine the effectiveness of mannose as a direct anti-fibrotic (Figure 4A). Mice were treated three times weekly with IP injections of 0.05 mL 20% CCl_4_ in corn oil (CO) or with 0.05 mL CO as the vehicle control for four weeks. CCl_4_-injected mice were given either standard drinking water or water supplemented with low (5%) or high dose (20%) mannose, either at the beginning of CCl_4_ administration as the “prevention” group, or after two-weeks delay as the “reversal” group. At 4 weeks, low and high dose mannose treatments in the prevention group reduced liver fibrosis as demonstrated by Sirius Red staining,*Col1a1* expression, and fibrosis scoring when compared to CCl_4_ mice with no mannose treatments (Figures 4B-E). Together, these data demonstrate that mannose supplementation has a novel role as an effective anti-fibrotic *in vivo,* both in a MASH and in a chemotoxic mouse model.

**Figure 4.**
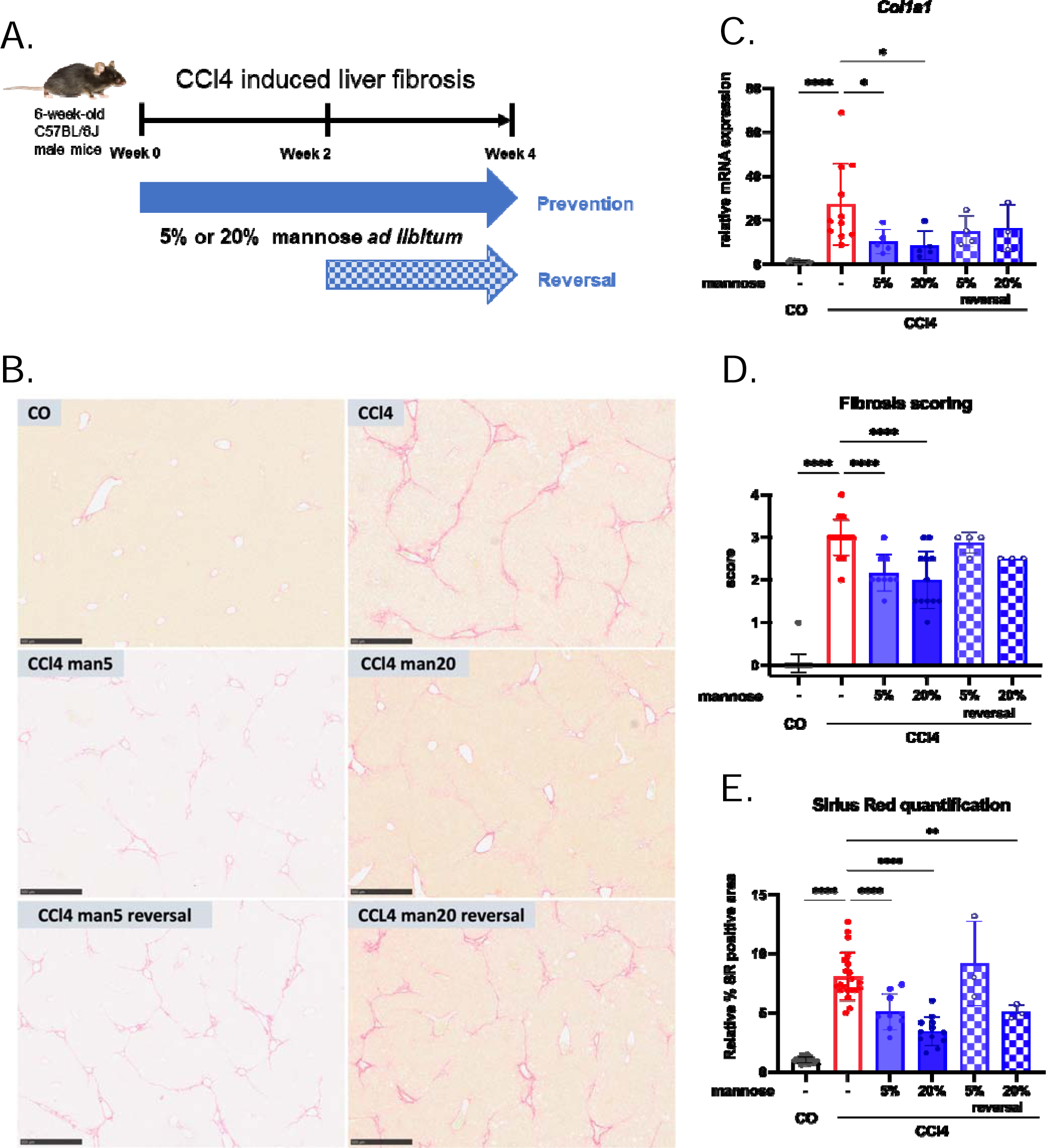
Mannose attenuates hepatic fibrosis in a CCl4 mouse model. (A) Graphical outline of the CCl_4_-induced liver fibrosis regimen. (B) Representative images of Sirius Red staining from whole liver sections. Scale bar: 500µm. (C) Quantitative PCR analysis of*Col1a1* mRNA expression; n=3-12. (D-E) Fibrosis scoring (D) and Sirius red quantification (E) quantified from Sirius red stained images; n=3-21. Results expressed as mean ± SD and compared by Student’s t-test (*p<0.05, **p<0.01, ***p<0.001, and ****p<0.0001).

### Mannose effects are dependent on fructose-mediated steatosis and KHK

To gain mechanistic insight into how mannose improves MASH, we performed bulk RNA sequencing (RNA-seq) on livers from mice on the FAT-MASH diet (MASH) or normal chow (ND) for 12 weeks, without or with mannose under the treatment protocols described (Figure 1A). Across all treatment groups, 83% of transcriptional variation was captured in the first 2 principal components (PC1: 75%, PC2: 8%, Figure 5A). Mice on the FAT-MASH regimen showed a strong transcriptional alteration compared to a normal diet (Figure 5A), confirming that the FAT-MASH diet induces robust transcriptional changes. In ND mice, low and high dose mannose had little, if any, effect on overall hepatic transcription. For FAT-MASH mice, mannose induced substantial transcriptional changes in a dose-dependent manner (Figure 5A). Surprisingly, principal component (PC) analysis revealed that the transcriptional changes did not revert to a “normal diet” state. Instead, changes appeared to be more complex, with transcriptional variance captured along PC1 expanding, whereas that along PC2 collapsing and reversing. Gene set enrichment analysis (GSEA) on differentially expressed genes was performed and compared across treatment groups (Supplemental Figure S2). Using hallmark gene set signatures, the ADIPOGENESIS and FATTY_ACID_METABOLISM pathways were among the most downregulated in MASH mice treated with mannose, compared to untreated MASH, and CHOLESTEROL_HOMEOSTASIS was upregulated in mannose-treated groups compared to the MASH group (Supplemental Figures S2 and S3).

**Figure 5.**
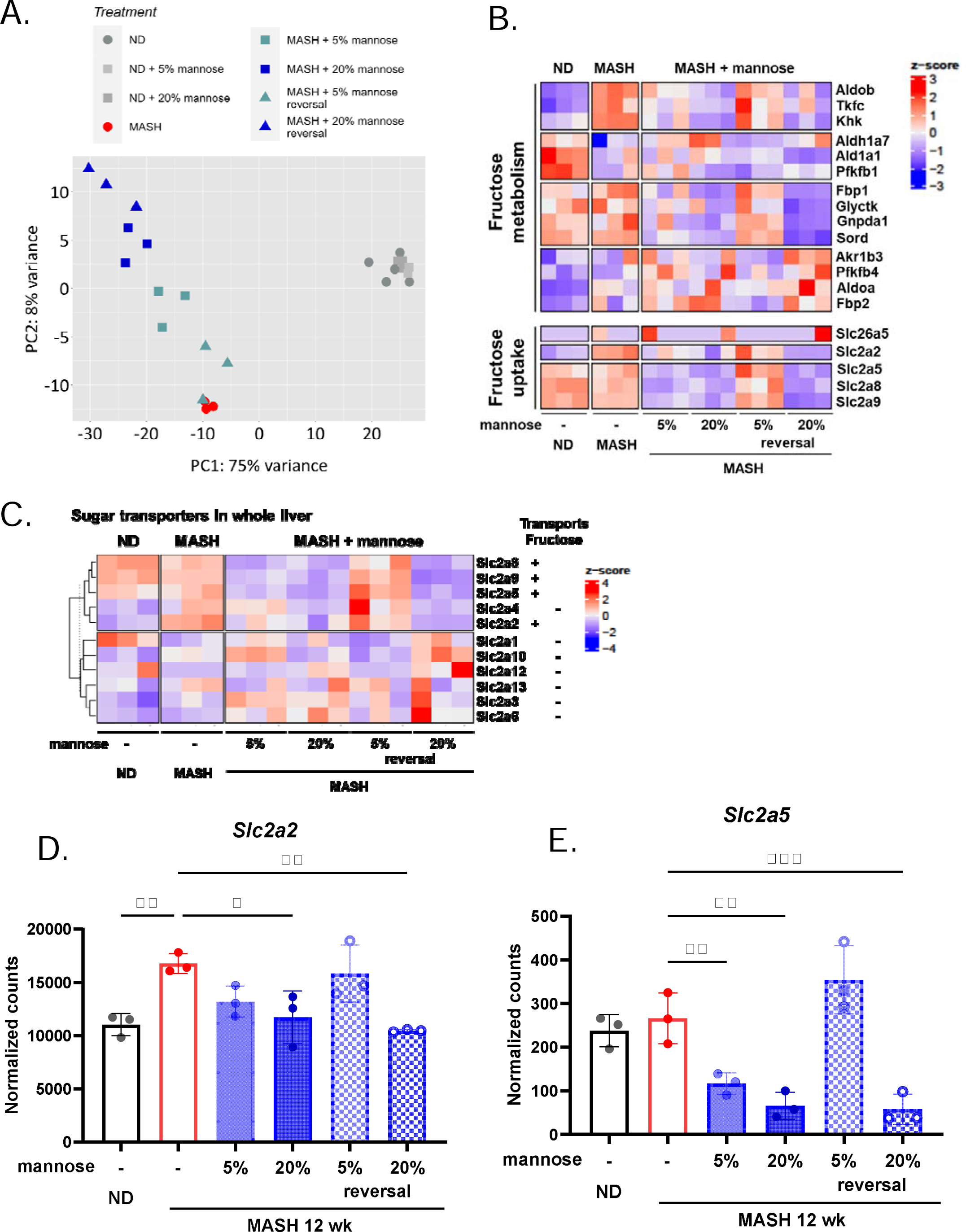
Bulk RNA-seq reveals that mannose induces transcriptional changes in fructose metabolism and uptake genes. (A) PCA plot of differentially expressed genes identified through bulk RNA sequencing of whole livers from ND and MASH livers without or with mannose supplements (n=3-5 per group). The first 2 principal components (83% of transcriptional variation across groups) are plotted. (B) Heat map showing normalized z-scores for expression of genes involved in fructose metabolism and uptake. (C) Heat map showing normalized z-scores for expression of genes involved in transmembrane sugar transport (*Slc2a* family). Transporters that are reported to transport fructose are indicated. (D-E) Bar plots showing normalized expression of fructose transporters *Slc2a2* encoding Glut2 (D) and *Slc2a5* encoding Glut5 (E) are shown. Statistical comparisons are by one-way ANOVA with Dunnett’s post hoc test for multiple comparisons (*p<0.05, **p<0.01, ***p<0.001).

Excessive fructose consumption is proposed as an important driver of MASLD and MASH^5–7,12^ and is a critical component of the FAT-MASH regimen to produce the MASH phenotype in mice. Mannose and fructose are metabolically linked through interconversion of their respective 6-phosphates (mannose 6-P ↔ fructose 6-P). Therefore, we hypothesized that mannose could exert its positive effects by modulating fructose metabolism. Indeed, the gene ontology analysis of liver RNA-seq data revealed that genes involved in fructose metabolism and fructose uptake were significantly changed in mannose-treated groups (Figure 5B). Genes encoding fructose transporters *Slc2a2*, *Slc2a5*, *Slc2a8*, and *Slc2a9* were downregulated by mannose (Figure 5C). These fructose transporters are members of the SLC2A (GLUT) family of facilitative hexose transporters, and we examined if mannose also modulated the expression of their other family members. Of the 14 known*Slc2a* family members, we found 11 were expressed in these liver samples. Except for*Slc2a4*, all of these fructose transporter genes were decreased in response to mannose, whereas those that do not transport fructose showed no expression changes (Figure 5C). This suggests that mannose specifically targets fructose uptake and thus, we specifically queried SLC2A2 and SLC2A5, which are the main fructose transporters. We found that mannose significantly dampened the expression of these transporters (Figure 5D-E).

For the fructose metabolism genes, the FAT-MASH diet increased expression of*Khk*, *AldoB*, and *Tkfc*, the first three steps of fructose metabolism, whereas mannose counteracted this increased expression (Figures 5B and 6A-B). Of note, the low dose mannose reversal group had minimal effect. For subsequent analyses, we focused on KHK as it is the first and rate-limiting step of fructose metabolism, by converting fructose to fructose-1-phosphate. KHK inhibition has demonstrated anti-steatotic effects in pre-clinical*in vitro* and *in vivo* models^17–19^, and investigations of KHK inhibitors have also reached the level of clinical trials for MASH (NCT03256526, NCT03969719, NCT05463575). By western blot in MASH liver, KHK was increased at 12-weeks livers compared to normal diet controls, and mannose reduced KHK expression in all mannose treated groups (Figure 6C). These data support that mannose supplementation dampens KHK and mitigates hepatic steatosis*in vivo*, and may offer an exciting novel therapeutic option as a KHK inhibitor.

**Figure 6.**
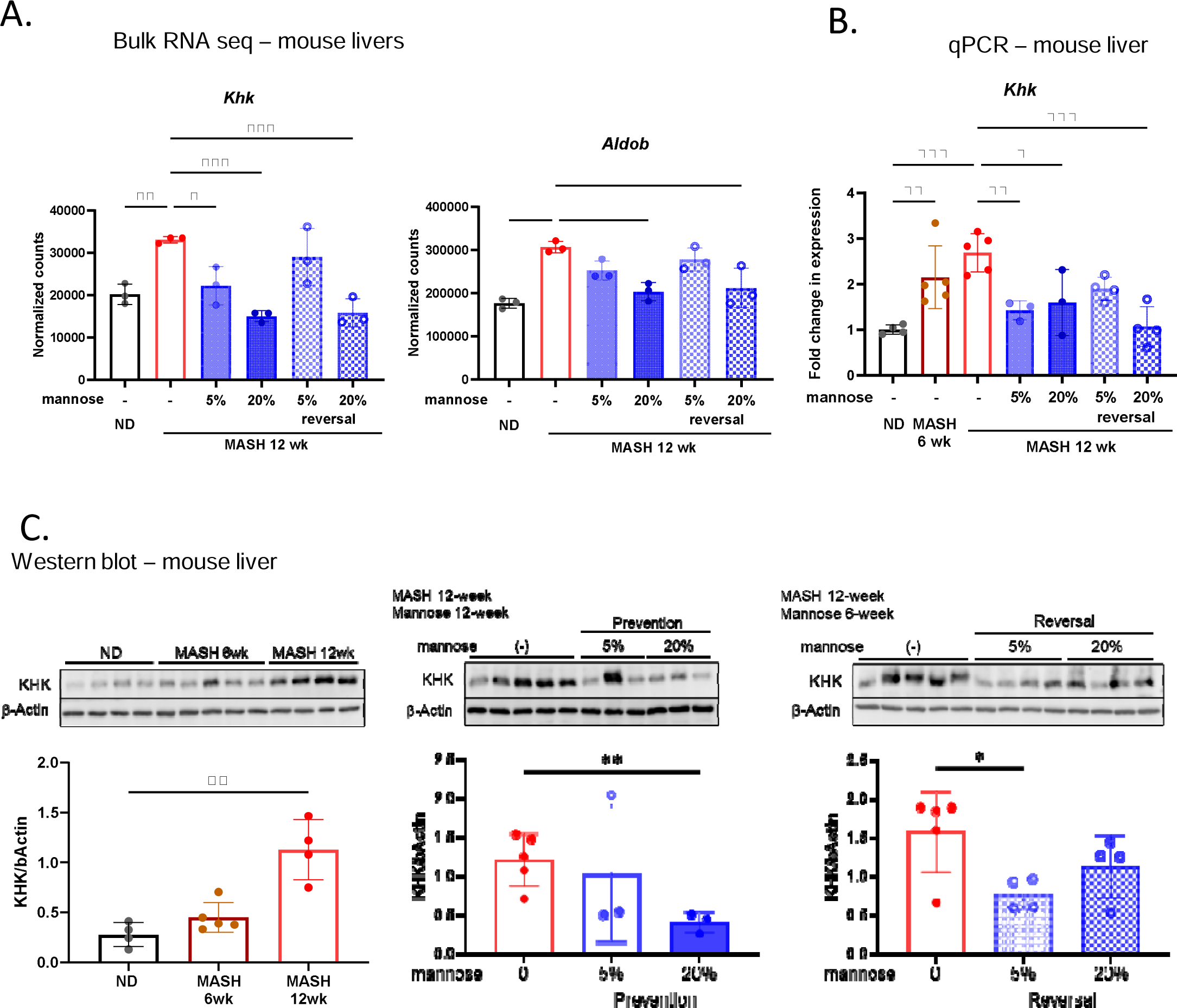
Mannose induces changes in expression of genes and proteins critical for fructose metabolism. (A) Bar plots showing normalized expression of*Khk* and *Aldob* identified by bulk RNA sequencing of whole liver transcriptomes from ND and MASH mice without or with mannose supplements. (B) Validation of *Khk* expression by qPCR. Statistical comparisons are by one-way ANOVA with Dunnett’s post hoc test for multiple comparisons (*p<0.05, **p<0.01, ***p<0.001). (C) Western blot analysis of KHK in whole liver lysates, withβ-Actin used for loading control. Semi-quantitative densitometry (ImageJ) of KHK/β-Actin ratios are shown, with comparisons by Student’s t-test (*p<0.05, **p<0.01).

Given our *in vivo* data showing that mannose dampened fructose metabolism pathways and KHK expression, we next examined the cell-specific role of mannose in hepatocytes and asked whether the anti-steatotic effects of mannose in hepatocytes were dependent on the presence of fructose. To test this, we mimicked the FAT-MASH conditions in primary mouse hepatocytes using a treatment protocol containing free fatty acids (FFA) and fructose^38,39^. Incubation of primary hepatocytes with 400 µM oleic + palmitic acid (2:1 cocktail) and 50 mM fructose for 72 hours induced steatosis as assessed by increased ORO staining (Figure 7A). Similar to findings in the FAT-MASH mice treated with mannose, primary mouse hepatocytes treated with mannose supplements demonstrated dampened steatosis, with 25 mM mannose treatment decreasing steatosis by 30% (Figure 7B). However, the anti-steatotic effects of mannose were dependent on the presence of fructose, as there was no improvement in steatosis in hepatocytes exposed to FFAs alone and treated with mannose (Figure 7A-B). Additionally, these findings were validated in the human hepatocyte cell line THLE-5B using 750 µM oleic + palmitic acid (2:1 cocktail) without or with 100 mM fructose for 72 hours^38,39^ and supplemented with either 10 or 25 mM mannose for 72 hours. Similar to the improvement in steatosis we observed *in vivo* and in primary mouse hepatocytes (Figures 1 and 7A-B), mannose treatments decreased steatosis by 33% in human hepatocytes (Figure 7C-D). However, this was again only seen in FFA + fructose conditioned hepatocytes; steatosis was unchanged when mannose was added to the cells conditioned with FFA alone (Figure 7C-D). These effects were specific to mannose since supplementing with galactose at the same concentrations failed to improve steatosis in FFA conditioned cells regardless of the presence of fructose (Supplemental Figure S4).

**Figure 7.**
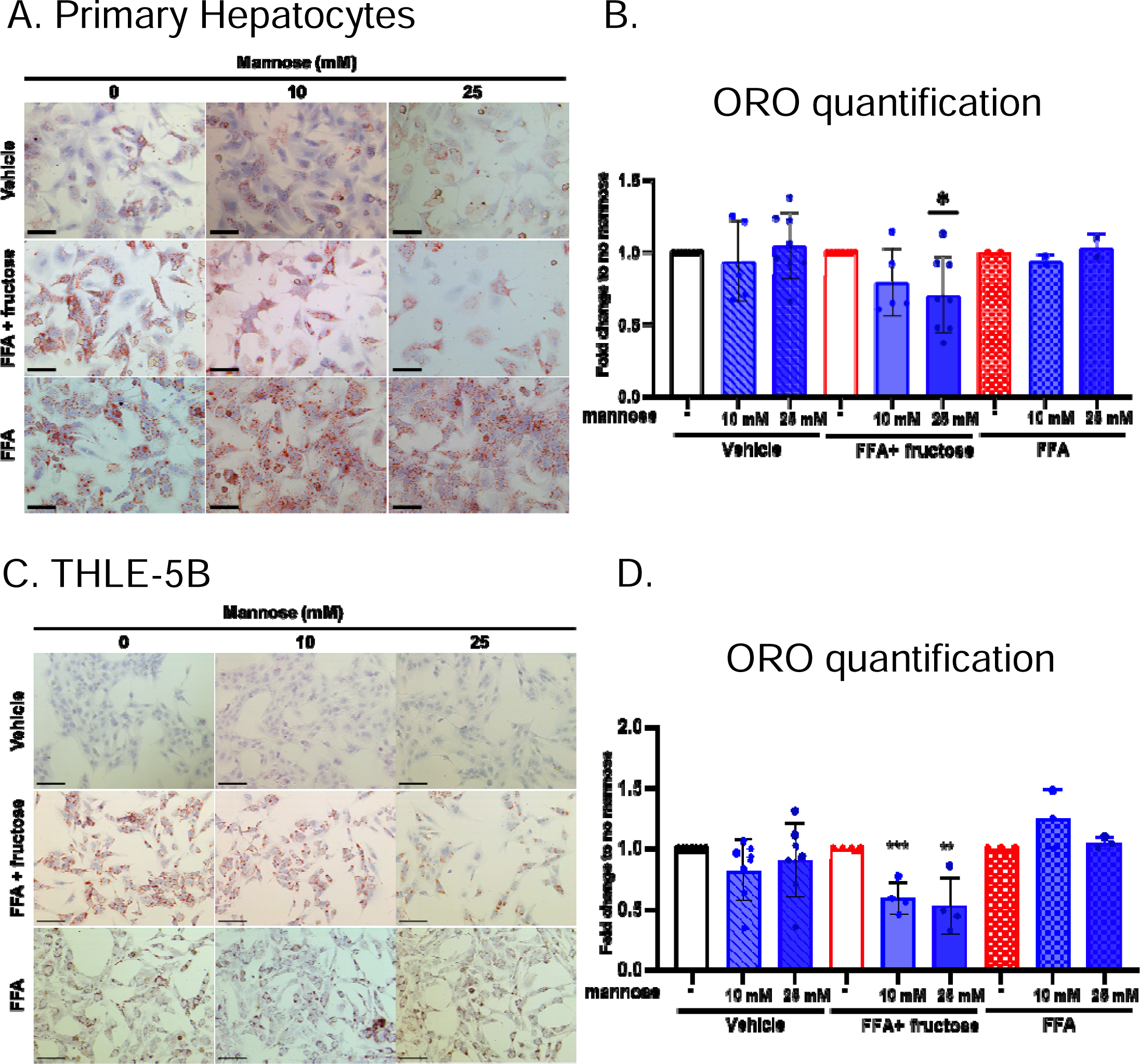
Mannose reduces steatosis *in vitro* and is fructose-mediated. (A, C) Photomicrographs of ORO-stained primary mouse hepatocytes (A) or THLE-5B human hepatocytes (C). Scale bar 100 um. (B, D) Bar charts of normalized ORO quantification in primary mouse hepatocytes (n=2-8, B) or THLE-5B human hepatocytes (n=3-7, D). Results are expressed as fold change to no mannose treatments in each condition and analyzed with one sample t-test (*p<0.05, **p<0.01, ***p<0.001).

We explored whether KHK was altered in primary mouse hepatocytes under MASH conditions without or with mannose treatment. We performed immunofluorescence (IF) of FFA + fructose treated primary mouse hepatocytes with and without 25 mM mannose supplementation. Similar to the FAT-MASH mouse livers (Figure 6C), primary mouse hepatocytes treated with FFA + fructose showed increased KHK by IF, and mannose supplementation dampened KHK expression (Figure 8A). Similarly, in THLE-5B human hepatocytes, KHK expression by western blot was increased in FFA + fructose and FFA groups; mannose supplementation dampened KHK, but only in the presence of fructose (Figure 8B). These findings support the conclusion that mannose improved steatosis by dampening KHK. Thus, we tested whether KHK overexpression would abrogate the anti-steatotic effects of mannose supplementation. To do this, we overexpressed KHK in THLE-5B human hepatocytes. KHK was successfully overexpressed as detected by WB (Figure 8C). In hepatocytes with KHK overexpression, mannose was unable to improve steatosis (Figure 8D). These data demonstrate a therapeutic role for mannose in fructose-induced hepatic steatosis by dampening KHK expression.

**Figure 8.**
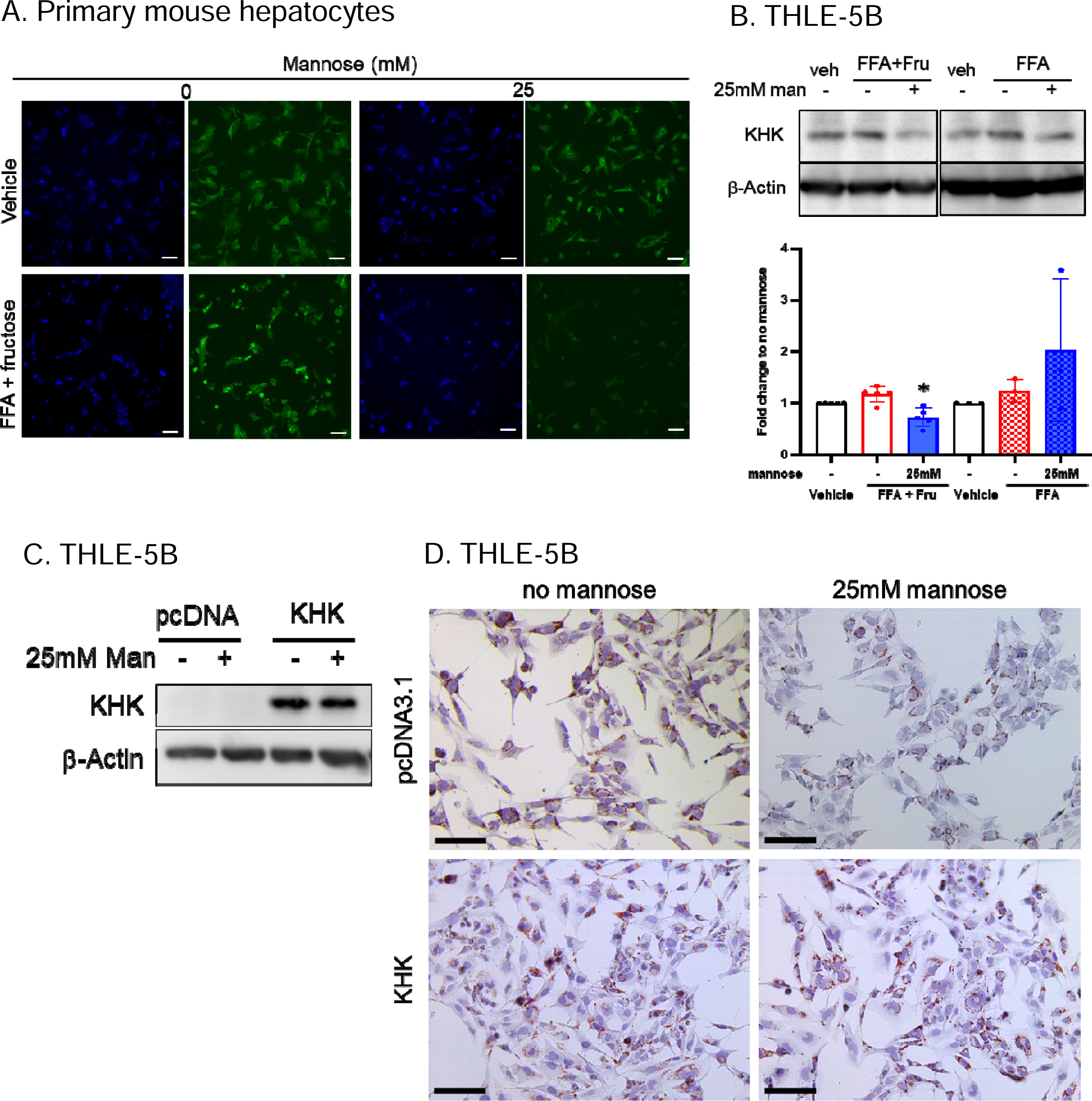
KHK is decreased in mannose-treated human hepatocytes (THLE-5B) and overexpression abrogates steatosis reduction. (A) Immunofluorescent staining for KHK (green) in primary mouse hepatocytes treated with vehicle or FFA + fructose, without or with mannose, for 72 hours. Cells are counterstained with DAPI (blue). Scale bar = 100 um. (B, upper) Representative western blot for KHK in THLE-5B human hepatocytes treated with vehicle, or FFA + fructose (FFA+Fru) or FFA alone, without or with 25 mM mannose.β-Actin is used as loading control. (B, lower) Bar chart showing semi-quantitative densitometry (ImageJ) of KHK/β-Actin normalized expression. Fold changes to no mannose treatments for each condition are shown (n=3-5), with comparisons by one sample t-test (*p<0.05). (C) Representative western blot showing overexpression of KHK or empty vector control (pcDNA3.1) in THLE-5B human hepatocytes, treated with FFA + fructose, without or with 25 mM mannose for 48 hours. β-Actin is used for loading control. (D) ORO staining of THLE-5B cells treated with FFA + fructose, without or with 25 mM mannose, and expressing endogenous KHK or empty vector (pcDNA3.1). Scale bar 100 um.

## Discussion

Mannose is a simple sugar, which is well-tolerated and readily accessible. Our study is the first to demonstrate that mannose supplementation has potential therapeutic roles in both MASH-induced steatosis and fibrosis. Here, we show that mannose supplementation in the drinking water of FAT-MASH mice can attenuate and reverse MASH-steatosis and fibrosis*in vivo.* In the FAT-MASH mice treated with mannose supplementation, H&E and ORO staining from the whole livers showed qualitative reduction in steatosis, decreased liver TG and cholesterol levels, and quantitative improvements in both steatosis and fibrosis using an unbiased AI-based platform (FibroNest^TM^).

We establish that the anti-steatotic effect of mannose is mediated by fructose. Therapeutic approaches that target fructose metabolism is of particular importance as fructose consumption in humans has been demonstrated to increase hepatic fatty acid synthesis^40^ and is associated with higher MASH fibrosis stage^41^. Clinical trials in adults^42^ and children^8^ demonstrate that isocaloric fructose restriction improves hepatic fat content. In our study, bulk RNA-seq from the FAT-MASH mouse livers demonstrated that mannose treatment dampened fructose metabolism and uptake genes. In both mouse and human hepatocytes, we demonstrated that mannose reduced steatosis, but only in fructose-conditioned hepatocytes (FFA + fructose) and not in the FFA alone condition. Our data points to KHK as the mechanism by which mannose regulates liver steatosis. This is supported by bulk liver RNA-seq of FAT-MASH mice, qPCR, and western blots of mouse livers in which KHK was increased in MASH conditions, and dampened with mannose supplementation. In primary mouse hepatocytes and in human hepatocytes, KHK protein expression is decreased in FFA + fructose conditions with mannose treatments, and this is not observed with FFA alone. Lastly, the anti-steatotic effect of mannose was abrogated when KHK was overexpressed in human hepatocytes.

Mannose may improve steatosis through decreased KHK by several mechanisms. First, it is possible that the decreased KHK expression leads to reduced fructolysis, which in turn leads to lower intracellular levels of acetyl CoA, the main substrate for DNL, and other downstream metabolites. Subsequently, the altered availability of necessary substrates for DNL leads to reduced steatosis. Another possible mechanism is through increasing expression of carnitine palmitoyltransferase 1A (CPT1A), a major mitochondrial beta-oxidation protein. Recently, it has been found that an increase in KHK expression leads to lowered CPT1A protein levels via acetylation at K508 in mice given high fat and fructose diet^13,43^ and mannose may be restoring CPT1A levels by reducing KHK expression. Lastly, mannose may reduce ER stress caused by increased KHK expression, which is implicated in the pathogenesis of MASLD in mouse models^13,43^.

Mannose may reduce steatosis through other components of fructose metabolism including: 1) decreasing fructose uptake, 2) reducing direct activation of DNL, and 3) mitigating intestinal inflammation and dysbiosis in the microbiome. Our bulk liver RNA-seq findings showed that fructose uptake pathways were altered, with lowered expression of fructose transporters, such as GLUT2, GLUT5, and GLUT8, when mice were given mannose. This suggests that mannose may be affecting fructose transporters. Given the structural similarity of mannose and fructose, competitive inhibition can be considered as well; this possibility could be tested with fructose transporter specific competitive inhibitors. GLUT8 is an interesting target as DeBosch et al. found that GLUT8 knockout mice were resistant to fructose-induced DNL and macrosteatosis^44^. Fructose induces hepatic DNL, and our bulk RNA-seq shows alterations in lipid homeostasis and lipogenesis. While the exact mechanisms are still being elucidated, key lipogenic regulator genes such as chREBP and SREBP1 are shown to be increased with fructose^45,46^; future investigations into the effects of mannose on these targets and on lipidomic profiling would be of interest. Lastly, it has been shown that fructose can cause direct intestinal epithelial damage^47^ and microbiome dysbiosis^48^, both of which result in gut inflammation via increased permeability, microbial/LPS translocation, and T cell modulation. These changes ultimately contribute to pathogenesis of MASLD. Mannose has been shown to induce T_reg_ cell immune response^22^, suppress macrophage IL-1b production^49^, and alter the microbiome to reduce obesity in mice given a high fat diet^21^. Recently, Dong et al. showed that mannose attenuated chemically-induced and spontaneous murine models of inflammatory bowel disease by protecting the integrity of the epithelial barrier^50^.

While mannose supplementation reduced MASH-induced steatosis and fibrosis, mannose likely exerts anti-fibrotic effects independent of MASLD injury. Our group has previously shown that mannose dampens HSC activation*in vivo* and *in vitro*^26^, and mannose has also been shown to reduce pulmonary and renal fibrosis in murine models^51,52^. In this study, our data demonstrate decreased fibrosis in two distinct *in vivo* models, suggesting mannose is not dampening MASH-fibrosis solely through ameliorating steatosis. Our data is based on whole liver and cultured hepatocytes, thus further investigations using single cell resolution or spatial techniques such as spatial transcriptomics are necessary to further delineate the independent effects of mannose on HSCs and hepatocytes in MASH and liver fibrosis.

The global prevalence of MASH and its rise as a leading indication for liver transplant provide strong impetus for research efforts to identify effective therapeutics. Mannose is a simple sugar that is well-tolerated, easily accessible, and already in clinical use for a rare pediatric disease^53^. Our study is the first to uncover mannose supplementation as an exciting, potential therapy that can not only improve metabolic dysfunction-associated steatosis, but offers a potential new candidate to treat MASH-fibrosis. We show that mannose can improve MASH by dampening fructose metabolism, which has been shown to be a potent driver of MASH fibrosis^41^. Given the proven safety and tolerability of mannose, it is feasible to envision next steps would be to test the effects of mannose supplementation in clinical trials for human MASH patients.

### Materials and Methods Mice

Six-week-old male C57BL/6J mice were purchased from Jackson Laboratory (Bar Harbor, ME). Three to five mice were housed in each cage and kept in a 12-hour dark – 12-hour light cycle environment. All procedures were performed according to protocols approved by the Animal Care and Use Committee of Icahn School of Medicine at Mount Sinai (IACUC-2015-0050).

### FAT-MASH model

Eight-week-old male mice were divided into two groups of control and MASH conditions. Control mice were fed a normal chow diet (LabDiet, Rodent diet 20, #5053) and normal RO water *ad libitum*. MASH mice were placed on high fat, high sugar western diet comprised of a 21.1% fat, 41% sucrose, and 1.25% cholesterol by weight chow (Teklad diets, TD. 120528) and a 23.1 g/L d-fructose (Sigma-Aldrich, F0127) and 18.9 g/L d-glucose (Sigma-Aldrich, G8270) sugar water for 12 weeks. MASH mice also received 0.2uL/g of body weight CCl_4_ (Sigma-Aldrich, 289116-100ML) intraperitoneal (IP) injections once per week (Tsuchida et al., 2018). Mice were treated with 5 or 20% (w/v) oral mannose supplemented in drinking water (Fisher, AAA1084, with administration beginning at week 0 (“preventative”) or 6 (“reversal”). Experimental groups were as follows: ND (n=4), ND and 5% mannose (ND 5% man, n=2), ND and 20% mannose (ND 20% man, n=3), MASH at week 12 (MASH 12-w) (n=9), MASH at week 6 (MASH-6w, n=5), MASH and 5% mannose (MASH 5% man, n=8) MASH and 5% mannose beginning week 6 (MASH-rev 5% man, n=4), MASH and 20% mannose (MASH 20% man, n=9), and MASH and 20% mannose beginning week 6 (MASH-rev 20% man, n=4). Mice were euthanized by CO_2_ and cardiac puncture at week 6 or 12 for liver and serum samples.

### CCl**_4_** model

Mice were treated with IP injections of 0.05 mL 20% CCl_4_ (Sigma-Aldrich, 289116-100ML) in corn oil three times per week for four weeks; injections of 0.05 mL corn oil served as the vehicle control. Mice were treated with 5 or 20% oral mannose supplemented in drinking water (Fisher, AAA1084) with administration beginning at week 0 or week 2. Mice were euthanized by CO_2_ and cardiac puncture at week 4 for liver tissue and serum samples.

### Histology

Formalin-fixed liver pieces were embedded in paraffin, sectioned, and stained with either hematoxylin and eosin (H&E) or Picrosirius Red (Abcam, ab150681) by the Mount Sinai Biorepository and Pathology CoRE. For Oil Red O (ORO) staining, fresh liver sections were embedded in O.C.T. Compound (Fisher, 23-730-571) and flash frozen in liquid nitrogen. 10 um frozen sections were cut and stained with ORO (Fisher, AAA1298914). For fibrosis scoring and Sirius Red quantification in CCl_4_ model, stained slides were blinded. Quantification was performed using ImageJ software.

### Lipid measurements

Liver TG was extracted as previously described^54^. Briefly, liver tissue is homogenized and put into chloroform and methanol mix. The mix is separated into two phases – upper phase and lower phase containing lipids. Lower phase is reconstituted with 2-propanol and undergoes series of extraction reactions to isolate TG. Liver TG is then measured using Infinity triglyceride kit (ThermoFisher Scientific, TR22421).

### Serum Analysis

Serum ALT, AST, total bilirubin, and direct bilirubin levels were measured using VRL Diagnostics.

### Digital Pathology and AI

Liver histological sections were stained with Picrosirius Red for collagens and imaged at 20X in white light with an Aperio Digital Pathology system. FibroNestTM, a cloud based, high-resolution, single object image analysis platform, was used to quantify the fibrosis phenotype for collagen content and structure features (12 traits), fiber morphometry (12 traits), and fibrosis architecture (7 traits to measures the organization of the fibers). Color normalization and standardization^33,55^ was performed to calibrate the images of the study. Each trait was quantified with 7 quantitative parameters (qFTs) to account for severity, distortion, and variance, resulting in a total of 448 qFTs. Fibers were also classified into fine (simple skeleton or low Node/Branch ratio) and Assembled (complex skeleton or high Node/Branch ratio). The qFT dataset was automatically (AI) surveyed to identify traits (principal qFTs) that would exhibit a significant (p<0.05) and meaningful (>20%) relative difference (group average) between the Control and MASH groups. Such principal qFT are normalized to the non-steatotic and parenchymal tissue area (PT-prefix) and assembled into a normalized Phenotypic Fibrosis Composite Score (PT-Ph-FCS) and displayed in the form of a heat chart (Figure 3C). The parenchymal tissue normalization accounts for the effect of steatosis vacuoles being replaced by functional hepatocytes and is not performed on the architectural qFTs to protect the unique traits of fibrosis architecture (e.g. septal bridges) from any anti-steatotic effect.

Similarly, macro-steatotic vacuoles are identified. The percent macro-steatotic area of non-fibrotic tissues is computed using Medium fat vacuoles (6 um < equivalent diameter < 18 um) and Large vacuoles (equivalent diameter > 18 um) or for both combined (“All”). The normalized counts of medium, large, and “all” vacuoles (count/mm^2^) are used as markers for generation and growth of steatosis vacuoles and the effect of the intervention on such clusters. Small steatosis vacuoles (equivalent diameter < 6 um) are not accounted for in the method, as the algorithms is confused (poor performance) by other forms of “white space” in and around the hepatocytes. The FibroNest method has been applied to this MASH model and has demonstrated its superiority to conventional histological staging^32,56,57^. It has also been applied to other MASH rodent models and biological systems and is proven to verify the antifibrotic effect of reference compounds^32^.

### Liver bulk RNA-sequencing

Bulk RNA-seq was carried out by a commercial vendor (Novogene). Briefly, total RNA was prepared from flash-frozen whole mouse livers. Messenger RNA was purified from total RNA using poly-T oligo-attached magnetic beads. After fragmentation, first strand cDNA was synthesized using random hexamer primers followed by second strand cDNA synthesis. The library was ready after end-repair, A-tailing, adapter ligation, size selection, amplification, and purification. After QC, quantified libraries were pooled and sequenced on a NovaSeq PE150 platform (Illumina). Sequencing quality was confirmed (Q30>=91%) and aligned to mouse reference genome (GRCm39) using STAR RNA-seq aligner^58^. Read counts were generated (featureCounts, subread package^59^) and analyzed using DESeq2 for differential gene expression^60^. Three biological replicates were utilized for RNA-seq analysis.

### RNA isolation and qPCR analysis

Frozen liver was homogenized and lysed in TRIzol (ThermoFisher, 15596018) using Bel-Art^TM^ Proculture Cordless Homogenizer (Fisher Scientific, 17000201), and RNA was purified following supplied TRIzol protocol. RNeasy mini kit (Qiagen, 74106) was also used. Briefly, frozen liver was first homogenized and lysed. The lysate was loaded into the RNeasy silica membrane and bound for further washes. RNA is then purified using DNase treatment and eluted.For reverse transcription, we used the SuperScript complementary DNA (cDNA) synthesis kit (Quantabio, Beverly, MA). qRT-PCR was performed using PerfeCTa SYBR Green Fast Mix (Quantabio).

Samples were run in triplicate on the Roche LightCycler 480 in a reaction volume of 10 µL. Gene expression levels were normalized to *Gapdh* using the comparative threshold cycle (ΔΔCt) method. Primer sequences are mouse *Col1a1*: forward: mCol1a1-F1q - GTCCCTGAAGTCAGCTGCATA, reverse: mCol1a1-R1q – TGGGACAGTCCAGTTCTTCAT; mouse*Khk*: forward: mKhk-F1q – ATTCTGCACGCCTACAGCTTC, reverse: mKhk-R1q – TACGGGAGCCATTGGAGTTG; mouse *Gapdh*: mGapdh-F1q – CAATGACCCCTTCATTGACC, reverse: mGapdh-R1q – GATCTCGCTCCTGGAAGATG.

### Gene Set Enrichment Analysis (GSEA)

GSEA (Broad Institute, UC San Diego)^61,62^, was performed on mouse treatment group comparisons, using the mouse hallmark signature gene sets (MH). Analysis was run following standard workflow as described in the GSEA-MSigDB documentation, using total normalized counts (DESeq2) as input. Subsequent normalized enrichment scores for desired treatment comparisons were compiled for presentation using the R heatmap package. Genesets used for fructose metabolism and uptake analyses were GO_BP_FRUCTOSE_METABOLIC_ PROCESS (accession GO:0006000) and GO_BP_FRUCTOSE_TRANSMEMBRANE_TRANSPORT (accession GO:0015755), respectively.

### Primary mouse hepatocyte isolation

Primary mouse hepatocytes were isolated from wild-type C57BL/6J male mice aged over 12 weeks via two-step collagenase perfusion, as previously described^63^. Briefly, mice were anesthetized with ketamine and xylazine (ketamine 100mg/kg, xylazine 10mg/kg; IP injection). The inferior vena cava was catharized with a 22-guage catheter (BD Precision, 305165) and infused with 50mL perfusion solution I (1x HBSS + 0.5 mM EGTA; Cytiva) followed by 100mL of perfusion solution II (Medium 199 with 20 mM HEPES + 1% BSA + 100 units/mL Penicilin-streptomycin + 10 µg/mL Gentamycin + Collagenase type IV 75 mg/100 mL; Worthington Biochemicals). Following cell dissociation, cells were filtered with 100 um mesh cell strainers, and washed twice with hepatocyte plating medium (HPM; Medium 199 + 10% FBS + 100 units/mL Penicilin-streptomycin + 10 µg/mL Gentamycin). Hepatocytes were further purified using Percoll (Cytiva, 17089109) gradient centrifugation: 700 rpm for 10 min. Cells were plated at 5×10^5^ cells/mL in HPM on rat tail type-1 collagen (Fisher, CB-40236) coated culture plates.

### Cell culture and KHK overexpression

THLE-5B human hepatocytes were maintained in complete medium (Dulbecco’s modified Eagle’s medium [DMEM] supplemented with 10% FBS, 2 mM L-glutamine, and 1x penicillin/streptomycin, Cytiva, SH30081.05), and routinely tested for mycoplasma using the Venor GeM Mycoplasma Detection Kit (Millipore-Sigma, MP0025). THLE-5B cells were plated in both 24 and 6-well plates and transiently transfected using Lipofectamine™ 3000 according to the manufacturer’s recommendations of 0.5 µg and 2.5 µg of DNA per well, respectively.

Human KHK driven by a CMV promoter (Plasmid pcDNA3.1-C-(k)DYK-KHK, GeneScript, OHu18645) was transfected into THLE-5B as well as pcDNA3.1-C-(k)DYK empty vector which was used as a control for transfections. Cells were allowed to settle overnight before subsequent transfection with either KHK or pcDNA3.1 constructs and then treated with various conditions of vehicle control and FFA + fructose without or with mannose 8 hours post-transfection. Cells were then collected for protein analysis or stained with ORO 48 hours after treatment and analyzed by western blot and light microscopy.

### *In vitro* MASH conditioning with free fatty acids and fructose

Oleic acid (Millipore Sigma, O1008) was prepared at 5 mM as a working stock by diluting in DMEM media with 1% lipid free BSA (Millipore Sigma, A8806). Palmitic acid (Millipore Sigma, P0500) was prepared at 5 mM as a working stock by warming to 70°C for 10 minutes and dissolving in 100% ethanol. Then, palmitic acid dissolved in ethanol was added to DMEM media with 1% lipid free BSA, which was then warmed to 55°C for 15 minutes, sonicated at room temperature for 10-15 minutes, and warmed again to 55°C for 15 minutes. Both oleic acid and palmitic acid were filter sterilized. Both working stocks contained 1% BSA and palmitic acid contained 1% ethanol. Oleic acid and palmitic acid working stocks were added to DMEM media at 2:1 ratio and further diluted to achieve the desired concentrations; 750 µM for THLE-5B human hepatocytes and 400 µM for primary mouse hepatocytes. By the end of oleic acid + palmitic acid condition preparation, treatments contained 0.2% ethanol. Therefore, vehicle controls contained 0.2% ethanol. Fructose (Sigma, F1027) was dissolved in DMEM media and filtered to achieve concentrations of 100 mM for the human hepatocytes and 50 mM for primary mouse hepatocytes. Hepatocytes underwent oleic acid + palmitic acid with and without fructose conditioning for 72 hours and the media was changed once at 24 hours.

### Mannose and galactose *in vitro* treatment protocol

Mannose was dissolved in either PBS or DMEM media to achieve the desired concentrations of 10 and 25 mM and filter sterilized. Mannose was given for 72 hours with one media change at 24 hours. Galactose treatment was done to confirm the specificity of mannose effects.

Galactose (Millipore Sigma, G0750) was dissolved in either PBS or DMEM media to achieve the desired concentrations of 10 and 25 mM and filter sterilized. Galactose was given for 72 hours with one media change at 24 hours.

### ORO staining and quantification

After undergoing the MASH conditioning and mannose treatment protocol above, hepatocytes were stained with ORO to measure steatosis. Cells were first washed with PBS, incubated with 10% formalin at room temperature for one hour, and washed with double-distilled water twice. Plates were incubated with 60% isopropanol for 5 minutes and dried completely. ORO was added for 10 minutes and washed with double-distilled water four to five times. Images were obtained at this time. ORO was eluted with 100% isopropanol. Optical density (OD) measurement for absorbance was taken at 500 nm using 100% isopropanol as a blank. For normalization, methylene blue staining was done. Methylene blue was destained with 40% methanol and 4% acetic acid. OD measurement was taken at 665 nm using 40% methanol and 4% acetic acid as a blank. ORO OD measurements were divided by corresponding methylene blue OD measurements to obtain normalized ORO OD measurements. Normalized ORO measurements were averaged according to the treatment group. Fold changes for normalized ORO OD measurements were calculated by dividing each condition treated with mannose over corresponding condition with no mannose.

### Protein preparation and western blot

Snap-frozen liver samples were thawed on ice and homogenized in RIPA lysis buffer (150 mM NaCl, 1% NP-40, 0.5% sodium deoxycholate, 0.1% SDS, 50 mM Tris, pH 7.4) + protease inhibitor cocktail (Roche Complete) using a Bel-Art ProCulture Cordless Homogenizer, followed by sonication using a Biorupter Plus cup sonicator (Diagenode). THLE-5B and primary hepatocyte cell cultures were scraped in PBS, centrifuged, and flash-frozen, followed by thawing on ice and sonication in RIPA lysis buffer + protease inhibitor cocktail. Protein lysates were quantified by BCA (Pierce, 23227), resolved by SDS-PAGE and electro-transferred onto PVDF membrane, blocked for 1 hour in 5% milk or 5% BSA in TBST, and incubated overnight in primary antibodies, followed by 1-2 hours incubation in HRP-conjugated secondary antibody, chemiluminescent detection using Pierce ECL substrate (Pierce, 32106) and imaged with a Bio-RAD ChemiDoc MP imaging system. Primary antibodies used were anti-KHK (ThermoFisher, PA5-29004; 1:1,000 in 2% milk/TBST), anti-Collagen-1 (Bioss, bs-10423R; 1:1,000 in 2% milk/TBST), anti-beta-Actin (GeneTex, GTX629630; 1:2,000 in 2% milk/TBST), and anti-alpha-Tubulin (DSHB, 12G10; 1:2,000 in 2% milk/TBST).

### Immunofluorescence and Microscopy

Cells were conditioned with 400 uM FFA + 50 mM fructose and treated with or without mannose for 72 hours on coverslips. Cells were then fixed with 4% paraformaldehyde in phosphate-buffered saline (PBS) for 30 mins at room temperature (RT). Cells were blocked with 0.1% BSA diluted in PBS with 0.2% saponin and 0.02% sodium azide, then subsequently labeled with primary anti-KHK antibody (Invitrogen, PA5-29004) diluted 1:100 in blocking solution for 1 hour and secondary antibodies also diluted in blocking solution for 45 min, all at RT. Coverslips were stained with 0.1mg/ml DAPI diluted in water (MP Biomedicals, 157574) and mounted onto glass slides using Vectashield Antifade Mounting Medium (Vector laboratories, H-1000) and imaged on a Leica DMi8 inverted LED fluorescent microscope. Species-or isotype-specific secondary antibodies from goat or donkey were conjugated to Alexa Fluor 488 (1:2000; Thermo Fisher Scientific). Analysis of fluorescent expression level and localization of anti-KHK was performed on acquired images using ImageJ/FIJI (1.54f, Java 1.8.0_322). Brightness and contrast were adjusted to visualize hepatocytes.

### Statistics

Data were analyzed using GraphPad Prism software (version 9.1.1; GraphPad Software Inc., San Diego, CA). Data are expressed as mean ± SD. Differences between experimental and control groups were analyzed by two-tailed, unpaired Student’s t-test or one-way analysis of variance (ANOVA), followed by Dunnet’s or Šidák post-hoc correction, when more than two groups were compared. Fold changes were analyzed by one sample t-test against theoretical mean of 1. *P*<0.05 was considered statistically significant.

## Acknowledgements

We thank Glenn Doherty and Deanna L. Benson at the Mount Sinai Microscopy and Advanced Bioimaging CoRE for assistance with Confocal and Lightsheet imaging; the Biorepository and Pathology CoRE for processing and staining of tissue samples at Mount Sinai;the computational and data resources and staff expertise provided by Scientific Computing and Data at the Icahn School of Medicine at Mount Sinai and supported by the Clinical and Translational Science Awards (CTSA) grant UL1TR004419 from the National Center for Advancing Translational Sciences.

## References

1. Rinella ME, Lazarus JV, Ratziu V, et al. A multisociety Delphi consensus statement on new fatty liver disease nomenclature. Hepatology. 2023;78(6):1966–1986.

2. Wong RJ, Aguilar M, Cheung R, et al. Nonalcoholic steatohepatitis is the second leading etiology of liver disease among adults awaiting liver transplantation in the United States. Gastroenterology. 2015;148(3):547–555.

3. Goldberg D, Ditah IC, Saeian K, et al. Changes in the Prevalence of Hepatitis C Virus Infection, Nonalcoholic Steatohepatitis, and Alcoholic Liver Disease Among Patients With Cirrhosis or Liver Failure on the Waitlist for Liver Transplantation. Gastroenterology. 2017;152(5):1090–1099.e1091.

4. Tacke F, Puengel T, Loomba R, Friedman SL. An integrated view of anti-inflammatory and antifibrotic targets for the treatment of NASH. J Hepatol. 2023;79(2):552–566.

5. Vos MB, Lavine JE. Dietary fructose in nonalcoholic fatty liver disease. Hepatology. 2013;57(6):2525–2531.

6. Jegatheesan P, De Bandt J-P. Fructose and NAFLD: The Multifaceted Aspects of Fructose Metabolism. Nutrients. 2017;9(3):230.

7. Jensen T, Abdelmalek MF, Sullivan S, et al. Fructose and sugar: A major mediator of non-alcoholic fatty liver disease. J Hepatol. 2018;68(5):1063–1075.

8. Schwarz J-M, Noworolski SM, Erkin-Cakmak A, et al. Effects of Dietary Fructose Restriction on Liver Fat, De Novo Lipogenesis, and Insulin Kinetics in Children With Obesity. Gastroenterology. 2017;153(3):743–752.

9. Schwimmer JB, Ugalde-Nicalo P, Welsh JA, et al. Effect of a Low Free Sugar Diet vs Usual Diet on Nonalcoholic Fatty Liver Disease in Adolescent Boys: A Randomized Clinical Trial. JAMA. 2019;321(3):256–265.

10. Cohen CC, Li KW, Alazraki AL, et al. Dietary sugar restriction reduces hepatic de novo lipogenesis in adolescent boys with fatty liver disease. J Clin Invest. 2021;131(24):e150996.

11. Lee D, Chiavaroli L, Ayoub-Charette S, et al. Important Food Sources of Fructose-Containing Sugars and Non-Alcoholic Fatty Liver Disease: A Systematic Review and Meta-Analysis of Controlled Trials. Nutrients. 2022;14(14):2846.

12. Herman MA, Birnbaum MJ. Molecular aspects of fructose metabolism and metabolic disease. Cell Metab. 2021;33(12):2329–2354.

13. Helsley RN, Park S-H, Vekaria HJ, et al. Ketohexokinase-C regulates global protein acetylation to decrease carnitine palmitoyltransferase 1a-mediated fatty acid oxidation. J Hepatol. 2023;79(1):25–42.

14. Park S-H, Helsley RN, Fadhul T, et al. Fructose induced KHK-C can increase ER stress independent of its effect on lipogenesis to drive liver disease in diet-induced and genetic models of NAFLD. Metabolism. 2023;145:155591.

15. Ouyang X, Cirillo P, Sautin Y, et al. Fructose consumption as a risk factor for non-alcoholic fatty liver disease. J Hepatol. 2008;48(6):993–999.

16. Softic S, Gupta MK, Wang G-X, et al. Divergent effects of glucose and fructose on hepatic lipogenesis and insulin signaling. J Clin Invest. 2017;127(11):4059–4074.

17. Gutierrez JA, Liu W, Perez S, et al. Pharmacologic inhibition of ketohexokinase prevents fructose-induced metabolic dysfunction. Mol Metab. 2021;48:101196.

18. Shepherd EL, Saborano R, Northall E, et al. Ketohexokinase inhibition improves NASH by reducing fructose-induced steatosis and fibrogenesis. JHEP Rep. 2021;3(2):100217.

19. Futatsugi K, Smith AC, Tu M, et al. Discovery of PF-06835919: A Potent Inhibitor of Ketohexokinase (KHK) for the Treatment of Metabolic Disorders Driven by the Overconsumption of Fructose. J Med Chem. 2020;63(22):13546–13560.

20. Porru D. Recurrent Urinary Tract Infections in Adult Women: a Pilot Study With Oral D Mannose. clinicaltrials.gov; 2014/04/25/ 2014. NCT01808755.

21. Sharma V, Smolin J, Nayak J, et al. Mannose Alters Gut Microbiome, Prevents Diet-Induced Obesity, and Improves Host Metabolism. Cell Rep. 2018;24(12):3087–3098.

22. Zhang D, Chia C, Jiao X, et al. D-mannose induces regulatory T cells and suppresses immunopathology. Nat Med. 2017;23(9):1036–1045.

23. Gonzalez PS, O’Prey J, Cardaci S, et al. Mannose impairs tumour growth and enhances chemotherapy. Nature. 2018;563(7733):719-723.

24. Zhang R, Yang Y, Dong W, et al. D-mannose facilitates immunotherapy and radiotherapy of triple-negative breast cancer via degradation of PD-L1. Proc Natl Acad Sci U S A. 2022;119(8):e2114851119.

25. Zhou X, Zheng Y, Sun W, et al. D-mannose alleviates osteoarthritis progression by inhibiting chondrocyte ferroptosis in a HIF-2α-dependent manner. Cell Prolif. 2021;54(11):e13134.

26. DeRossi C, Bambino K, Morrison J, et al. Mannose Phosphate Isomerase and Mannose Regulate Hepatic Stellate Cell Activation and Fibrosis in Zebrafish and Humans. Hepatology. 2019;70(6):2107–2122.

27. Hu M, Chen Y, Deng F, et al. D-Mannose Regulates Hepatocyte Lipid Metabolism via PI3K/Akt/mTOR Signaling Pathway and Ameliorates Hepatic Steatosis in Alcoholic Liver Disease. Frontiers in Immunology. 2022;13.

28. Fang J, Celton-Morizur S, Desdouets C. NAFLD-Related HCC: Focus on the Latest Relevant Preclinical Models. Cancers (Basel). 2023;15(14):3723.

29. Jiang M, Wu N, Chen X, et al. Pathogenesis of and major animal models used for nonalcoholic fatty liver disease. J Int Med Res. 2019;47(4):1453–1466.

30. Friedman SL, Neuschwander-Tetri BA, Rinella M, Sanyal AJ. Mechanisms of NAFLD development and therapeutic strategies. Nat Med. 2018;24(7):908–922.

31. Tsuchida T, Lee YA, Fujiwara N, et al. A simple diet-and chemical-induced murine NASH model with rapid progression of steatohepatitis, fibrosis and liver cancer. J Hepatol. 2018;69(2):385–395.

32. Wang S, Li K, Pickholz E, et al. An autocrine signaling circuit in hepatic stellate cells underlies advanced fibrosis in non-alcoholic steatohepatitis. Sci Transl Med. 2023;15(677):eadd3949.

33. Zheng Y, Jiang Z, Zhang H, Xie F, Shi J, Xue C. Adaptive color deconvolution for histological WSI normalization. Comput Methods Programs Biomed. 2019;170:107–120.

34. Carter JK, Bhattacharya D, Borgerding JN, Fiel MI, Faith JJ, Friedman SL. Modeling dysbiosis of human NASH in mice: Loss of gut microbiome diversity and overgrowth of Erysipelotrichales. PLoS One. 2021;16(1):e0244763.

35. Taylor RS, Taylor RJ, Bayliss S, et al. Association Between Fibrosis Stage and Outcomes of Patients With Nonalcoholic Fatty Liver Disease: A Systematic Review and Meta-Analysis. Gastroenterology. 2020;158(6):1611–1625.e1612.

36. Scholten D, Trebicka J, Liedtke C, Weiskirchen R. The carbon tetrachloride model in mice. Lab Anim. 2015;49(1 Suppl):4–11.

37. Fujii T, Fuchs BC, Yamada S, et al. Mouse model of carbon tetrachloride induced liver fibrosis: Histopathological changes and expression of CD133 and epidermal growth factor. BMC Gastroenterology. 2010;10(1):79.

38. Chen X, Li L, Liu X, et al. Oleic acid protects saturated fatty acid mediated lipotoxicity in hepatocytes and rat of non-alcoholic steatohepatitis. Life Sci. 2018;203:291–304.

39. Ramos MJ, Bandiera L, Menolascina F, Fallowfield JA. In vitro models for non-alcoholic fatty liver disease: Emerging platforms and their applications. iScience. 2022;25(1):103549.

40. Hochuli M, Aeberli I, Weiss A, et al. Sugar-sweetened beverages with moderate amounts of fructose, but not sucrose, induce Fatty Acid synthesis in healthy young men: a randomized crossover study. J Clin Endocrinol Metab. 2014;99(6):2164–2172.

41. Abdelmalek MF, Suzuki A, Guy C, et al. Increased fructose consumption is associated with fibrosis severity in patients with nonalcoholic fatty liver disease. Hepatology. 2010;51(6):1961–1971.

42. Simons N, Veeraiah P, Simons P, et al. Effects of fructose restriction on liver steatosis (FRUITLESS); a double-blind randomized controlled trial. Am J Clin Nutr. 2021;113(2):391–400.

43. Softic S, Meyer JG, Wang G-X, et al. Dietary Sugars Alter Hepatic Fatty Acid Oxidation via Transcriptional and Post-translational Modifications of Mitochondrial Proteins. Cell Metab. 2019;30(4):735–753.e734.

44. DeBosch BJ, Chen Z, Saben JL, Finck BN, Moley KH. Glucose transporter 8 (GLUT8) mediates fructose-induced de novo lipogenesis and macrosteatosis. J Biol Chem. 2014;289(16):10989–10998.

45. Ortega-Prieto P, Postic C. Carbohydrate Sensing Through the Transcription Factor ChREBP. Front Genet. 2019;10:472.

46. Kim M-S, Krawczyk SA, Doridot L, et al. ChREBP regulates fructose-induced glucose production independently of insulin signaling. J Clin Invest. 2016;126(11):4372–4386.

47. Todoric J, Di Caro G, Reibe S, et al. Fructose stimulated de novo lipogenesis is promoted by inflammation. Nat Metab. 2020;2(10):1034–1045.

48. Zhou X, Zhang X, Niu D, et al. Gut microbiota induces hepatic steatosis by modulating the T cells balance in high fructose diet mice. Sci Rep. 2023;13(1):6701.

49. Torretta S, Scagliola A, Ricci L, et al. D-mannose suppresses macrophage IL-1β production. Nat Commun. 2020;11:6343.

50. Dong L, Xie J, Wang Y, et al. Mannose ameliorates experimental colitis by protecting intestinal barrier integrity. Nat Commun. 2022;13(1):4804.

51. López-Guisa JM, Cai X, Collins SJ, et al. Mannose receptor 2 attenuates renal fibrosis. J Am Soc Nephrol. 2012;23(2):236–251.

52. Yamashita M, Niisato M, Kawasaki Y, Maemondo M. Protective roles of mannose receptor in an experimental pulmonary fibrosis model. European Respiratory Journal. 2018;52(suppl 62).

53. Niehues R, Hasilik M, Alton G, et al. Carbohydrate-deficient glycoprotein syndrome type Ib. Phosphomannose isomerase deficiency and mannose therapy. J Clin Invest. 1998;101(7):1414–1420.

54. Liu Q, Yu J, Wang L, et al. Inhibition of PU.1 ameliorates metabolic dysfunction and non-alcoholic steatohepatitis. J Hepatol. 2020;73(2):361–370.

55. Perez-Bueno F, Vega M, Sales MA, et al. Blind color deconvolution, normalization, and classification of histological images using general super Gaussian priors and Bayesian inference. Comput Methods Programs Biomed. 2021;211:106453.

56. Inia JA, Stokman G, Morrison MC, et al. Semaglutide Has Beneficial Effects on Non-Alcoholic Steatohepatitis in Ldlr-/-.Leiden Mice. Int J Mol Sci. 2023;24(10).

57. Ratziu V, Hompesch M, Petitjean M, et al. Artificial intelligence-assisted digital pathology for non-alcoholic steatohepatitis: current status and future directions. J Hepatol. 2023.

58. Dobin A, Davis CA, Schlesinger F, et al. STAR: ultrafast universal RNA-seq aligner. Bioinformatics. 2013;29(1):15–21.

59. Liao Y, Smyth GK, Shi W. featureCounts: an efficient general purpose program for assigning sequence reads to genomic features. Bioinformatics. 2014;30(7):923–930.

60. Love MI, Huber W, Anders S. Moderated estimation of fold change and dispersion for RNA-seq data with DESeq2. Genome Biol. 2014;15(12):550.

61. Subramanian A, Tamayo P, Mootha VK, et al. Gene set enrichment analysis: a knowledge-based approach for interpreting genome-wide expression profiles. Proc Natl Acad Sci U S A. 2005;102(43):15545–15550.

62. Mootha VK, Lindgren CM, Eriksson KF, et al. PGC-1alpha-responsive genes involved in oxidative phosphorylation are coordinately downregulated in human diabetes. Nat Genet. 2003;34(3):267–273.

63. Wang L, Yu J, Zhou Q, et al. TOX4, an insulin receptor-independent regulator of hepatic glucose production, is activated in diabetic liver. Cell Metab. 2022;34(1):158–170.e155.

