## Supplemental figures for "Mannose Supplementation Curbs Liver Steatosis and Fibrosis in Murine MASH by Inhibiting Fructose Metabolism"

### Slide 1
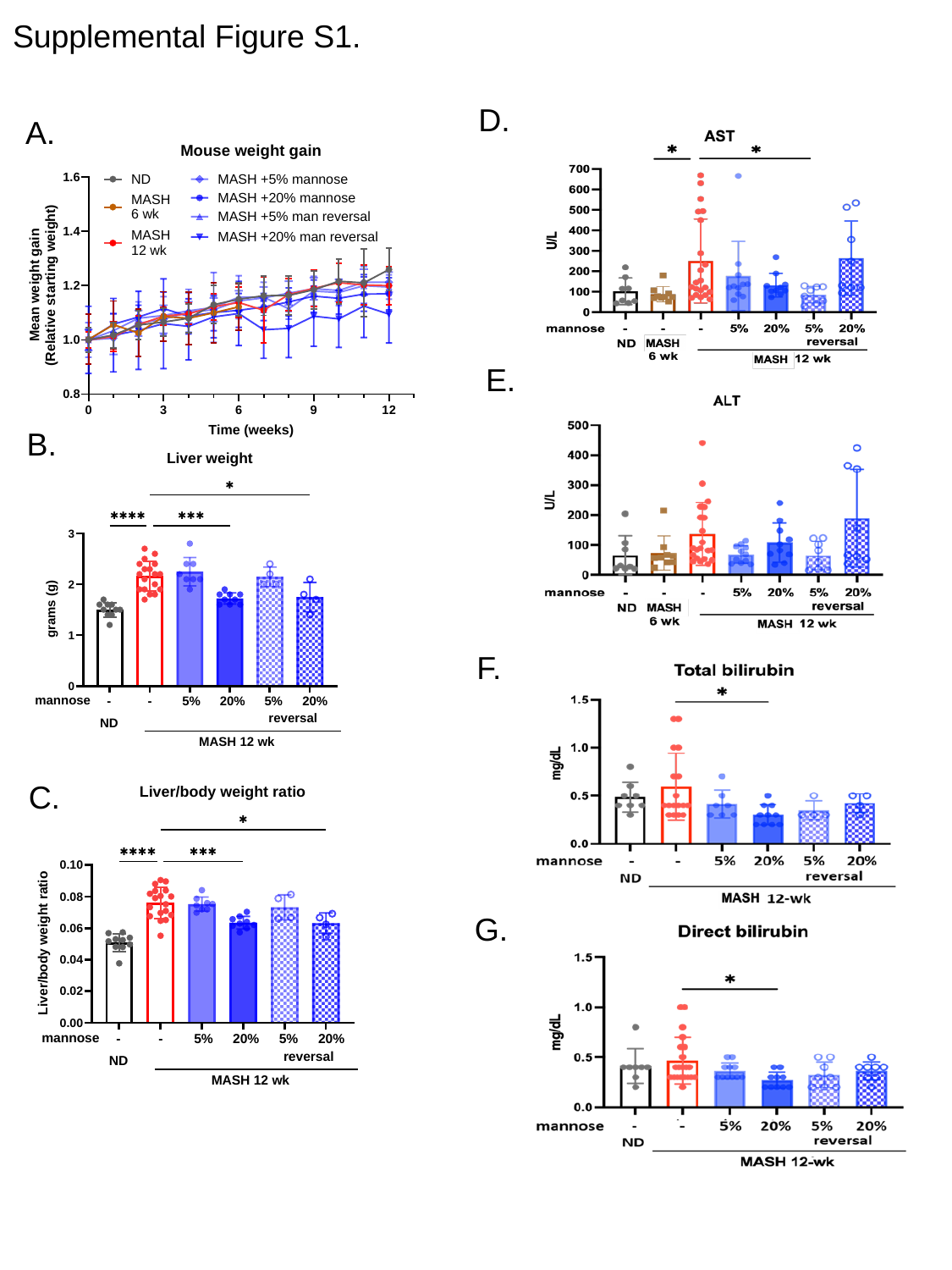

#### Supplemental Figure S1.
D.
A.
E.
B.
F.
C.
G.

### Slide 2
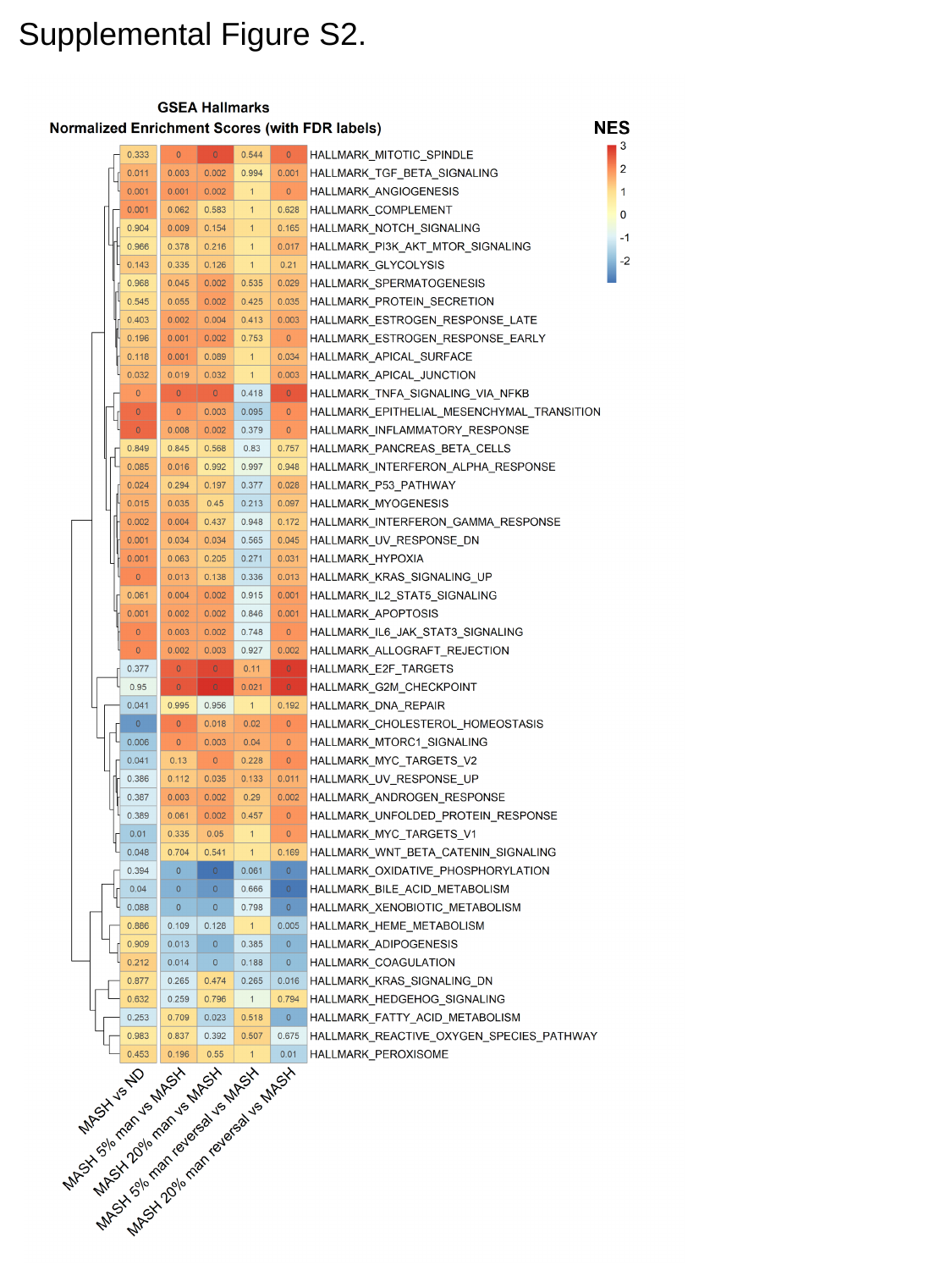

Supplemental Figure S2.

### Slide 3
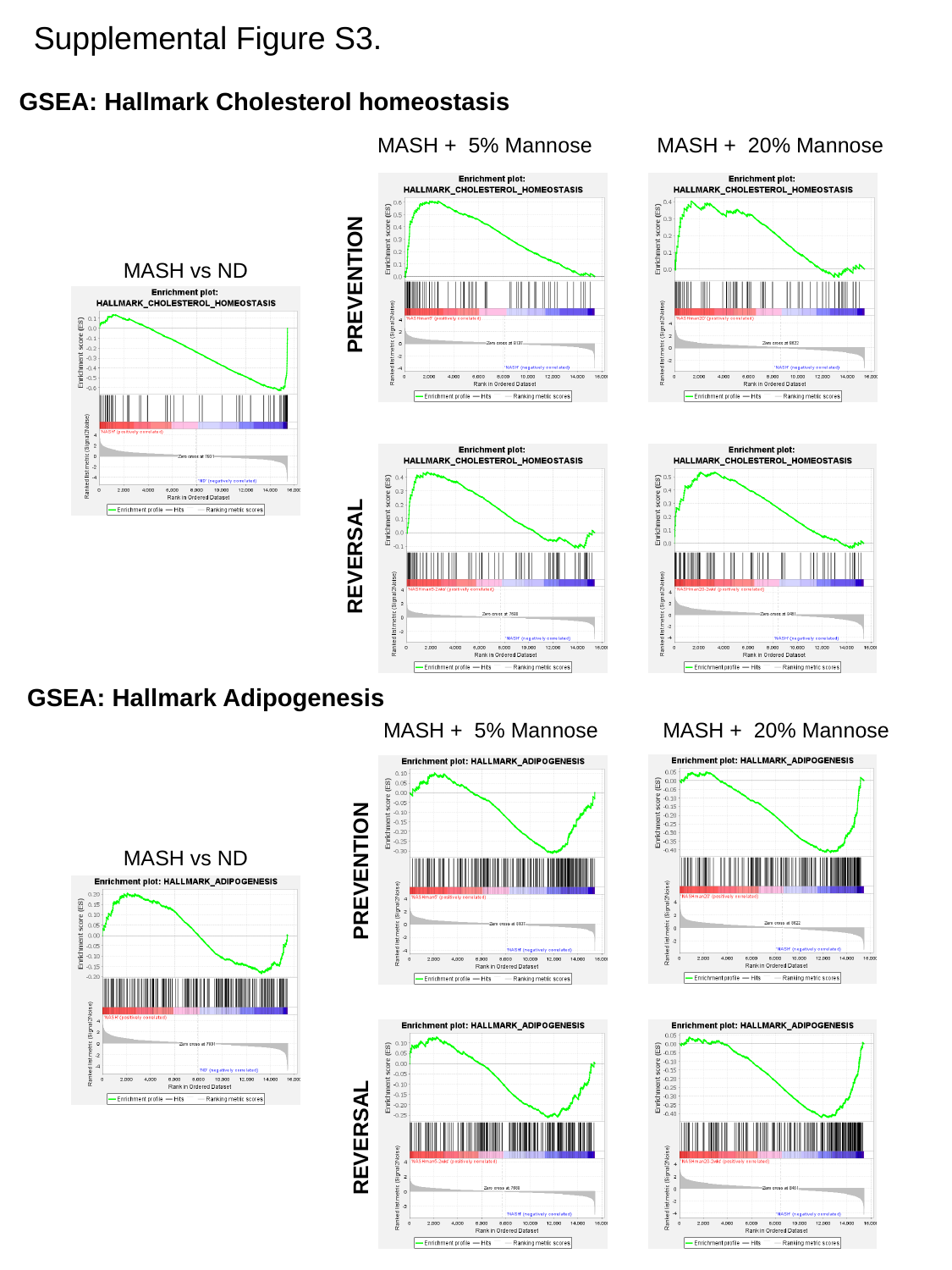

Supplemental Figure S3.
GSEA: Hallmark Cholesterol homeostasis
MASH + 5% Mannose
MASH + 20% Mannose
MASH vs ND
PREVENTION
REVERSAL
GSEA: Hallmark Adipogenesis
MASH + 5% Mannose
MASH + 20% Mannose
MASH vs ND
PREVENTION
REVERSAL

### Slide 4
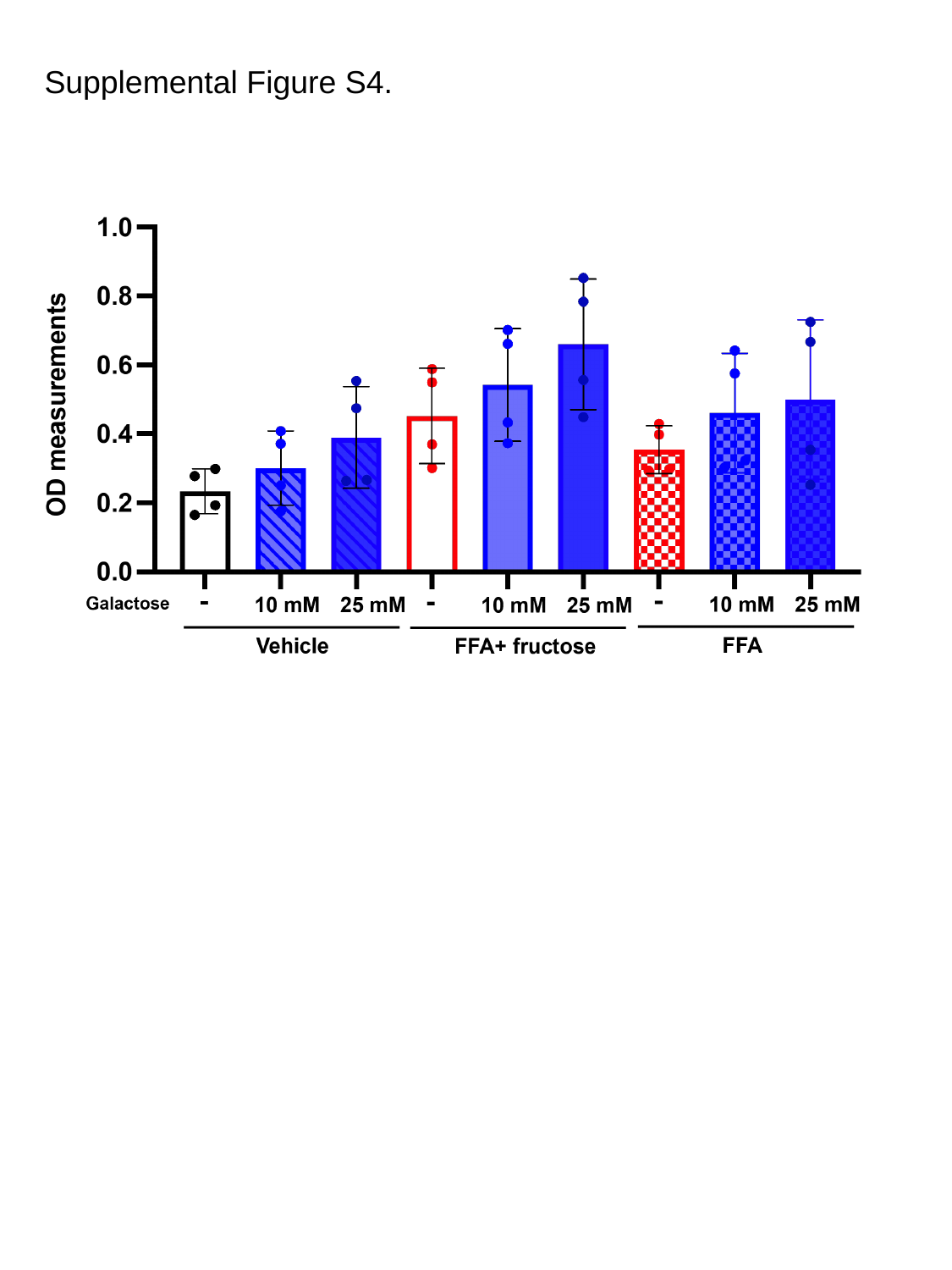

Supplemental Figure S4.

### Slide 5
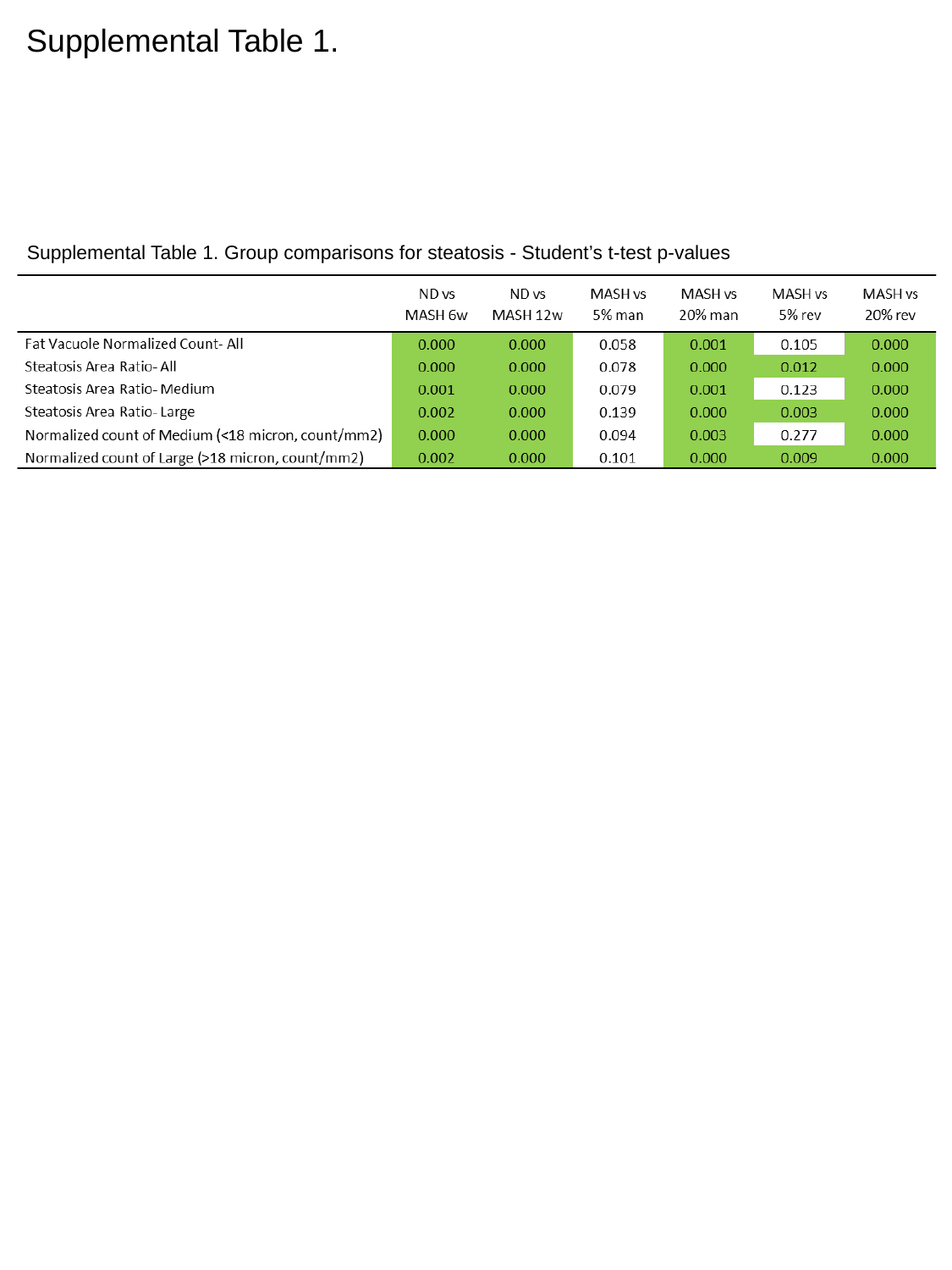

Supplemental Table 1.
Supplemental Table 1. Group comparisons for steatosis - Student’s t-test p-values

### Slide 6
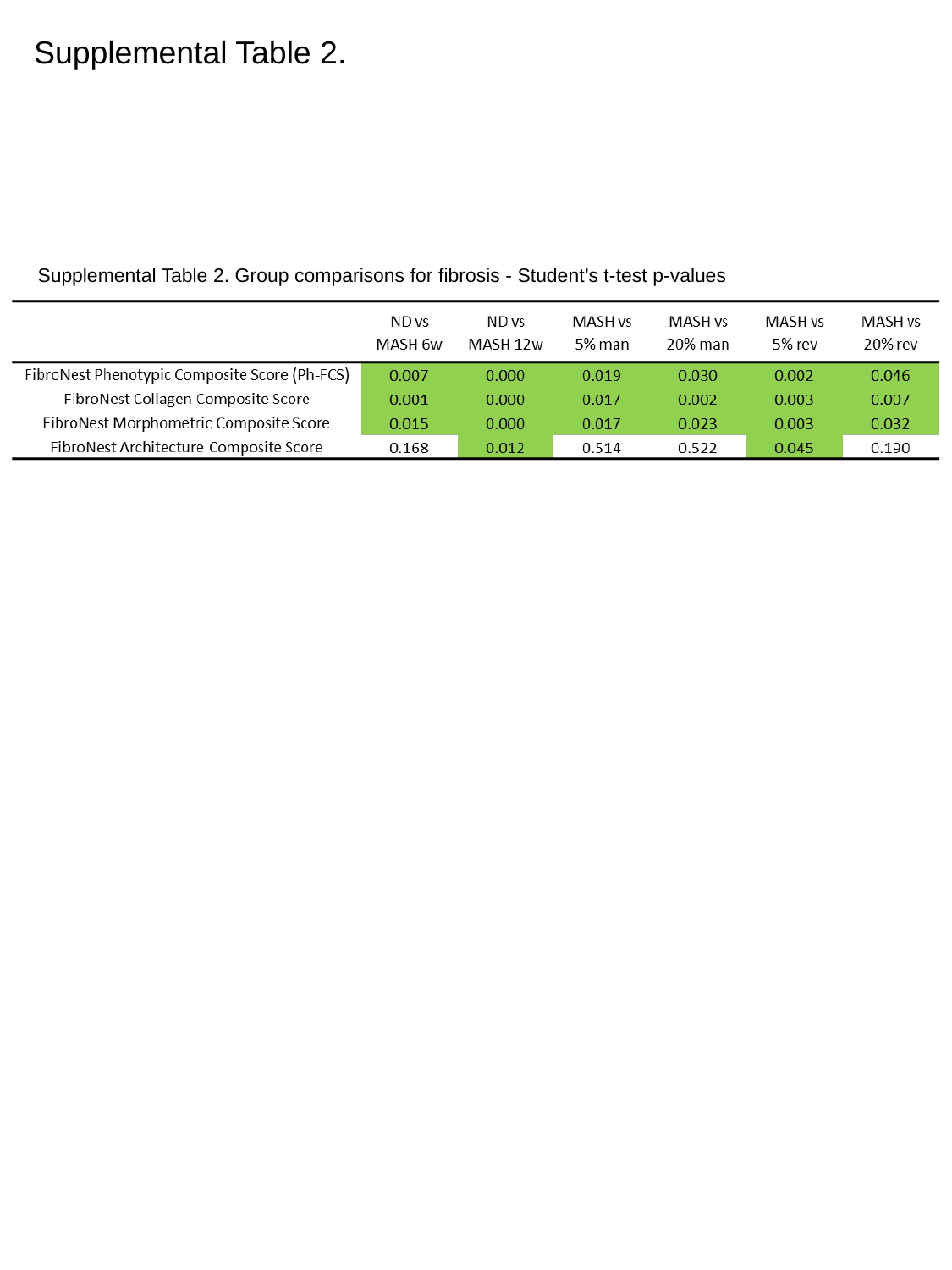

Supplemental Table 2.
Supplemental Table 2. Group comparisons for fibrosis - Student’s t-test p-values
