## Supplemental figure legends for "Mannose Supplementation Curbs Liver Steatosis and Fibrosis in Murine MASH by Inhibiting Fructose Metabolism"

**Supplemental Figure S1. Metabolic profile of FAT-MASH mice supplemented with mannose.** (A) Mouse total body weight measurements during course of MASH diet and mannose treatment regimens. Liver weight (B), liver to body weight ratio (C), and serum ALT (D), AST (E), total bilirubin (F), and direct bilirubin (G) measured at 12 weeks; n= 4-18 per group. Results are expressed as mean ± SD and were compared by Student’s t-test (*p<0.05, **p<0.01, ***p<0.001, and ****p<0.0001). ND, normal diet; MASH, FAT-MASH diet; man, mannose

**Supplemental Figure S2. Gene Set Enrichment Analysis (GSEA) for Hallmark pathways.** Heatmap of compiled results of GSEA from comparative gene expression patterns for different treatment groups of ND, MASH, and MASH + mannose. Hallmark gene set signatures were used for pathway identification. Normalized enrichment scores (NES) are indicated by heatmap color, and false discovery rate (FDR) are labeled within each cell.

**Supplemental Figure S3. Select differentially regulated GSEA Hallmark pathways.** Enrichment plots for select hallmark gene sets significantly enriched between indicated treatment groups.

**Supplemental Figure S4.** Bar plot showing THLE-5B hepatocytes conditioned with vehicle, FFA + fructose, or FFA alone, without or with 10 or 25 mM galactose for 72 hours (n=4).
